# Conserved Noncoding Cis-Elements Associated with Hibernation Modulate Metabolic and Behavioral Adaptations in Mice

**DOI:** 10.1101/2024.06.26.600851

**Authors:** Susan Steinwand, Cornelia Stacher Hörndli, Elliott Ferris, Jared Emery, Josue D. Gonzalez Murcia, Adriana Cristina Rodriguez, Tyler C. Leydsman, Amandine Chaix, Alun Thomas, Crystal Davey, Christopher Gregg

## Abstract

Our study elucidates functional roles for conserved *cis-*elements associated with the evolution of mammalian hibernation. Genomic analyses found topologically associated domains (TADs) that disproportionately accumulated convergent genomic changes in hibernators, including the TAD for the *Fat Mass & Obesity* (*Fto*) locus. Some hibernation-linked *cis-*elements in this TAD form regulatory contacts with multiple neighboring genes. Knockout mice for these *cis-*elements exhibit *Fto, Irx3,* and *Irx5* gene expression changes, impacting hundreds of genes downstream. Profiles of pre-torpor, torpor, and post-torpor phenotypes found distinct roles for each *cis-*element in metabolic control, while a high caloric diet uncovered different obesogenic effects. One *cis-*element promoting a lean phenotype influences foraging behaviors throughout life, affecting specific behavioral sequences. Thus, convergent evolution in hibernators pinpoints functional genetic mechanisms of mammalian metabolic control.

**One-sentence summary:** Deletions of conserved *cis-*elements tied to hibernator evolution causes diverse metabolic traits in mice.

---

Elucidating the genetic determinants of metabolic regulation is pivotal for understanding many key phenotypes in nature and medicine, including disease and aging processes (*1–4*). Species differences in metabolic traits could be leveraged to help uncover the genetics of metabolism. Hibernation constitutes an extreme metabolic phenotype (*5*, *6*). Obligate hibernators increase body weight by up to 30%–50%, and then enter torpor to survive months of food scarcity, suppressing metabolism and body temperature (*5–9*). In contrast, non-hibernating homeotherms maintain a relatively stable metabolic rate and internal body temperature throughout the year and forage using diverse strategies to survive variable seasonal conditions and food scarcity (*10*).

Studies of hibernator adaptations suggest that changing components of mammalian metabolism offers benefits such as energy balance control, neuroprotection (*11*), reversal of neurodegenerative processes (*12–14*), reversal of obesity and insulin resistance (*8*, *15*, *16*), and enhanced longevity (*17*). Uncovering the genetic mechanisms involved is expected to provide important advances in our understanding of mammalian metabolism and health.

Research on the genetic underpinnings of varied animal morphologies (*18–21*), as well as human genome-wide association studies (GWAS) (*22*), show that *cis-*regulatory mechanisms are key drivers of phenotypic diversity (*23*). However, while reporter assays (*24*) and other screens (*25*, *26*) can help find *cis-* regulatory elements (CREs) sufficient to affect gene expression, defining and demonstrating causal CREs and gene regulatory programs that govern complex metabolic, aging, and feeding behavior phenotypes remains daunting. Our companion study shows that convergent genomic changes in obligate hibernators illuminate specific CREs and cellular *cis-*regulatory programs governing molecular responses to fed, fasted, and refed (FFR) states in the mouse hypothalamus (*27*). Here, we present the functional characterization of these ‘hibernation-linked CREs’ using knockout mice, focusing on a subset that we show are components of a *cis-* regulatory program governing gene expression within the topological associated domain (TAD) containing the Fat Mass & Obesity (*Fto*) gene locus. TADs are 3-D genomic structures that function to limit CRE interactions to defined groups of neighboring genes and help delineate *cis-*regulatory programs in genomic regions (*28*). The *Fto* locus is the strongest genetic risk factor for human obesity (*29–31*), *Fto* knockout mice show reduced fat mass and abnormal growth (*32*, *33*), and the *Fto* TAD is a top region enriched for convergent genomic changes in hibernators based on our previous work (*28*). Our findings here show that individual hibernation-linked *cis-* elements are functional and affect varied aspects of gene expression, hibernation-relevant metabolic traits, and naturalistic foraging across the lifespan.

## The *Fto-Irx* Topologically Associated Domain is a Primary Genomic Region of Evolutionary Divergence between Hibernating and Homeothermic Mammals

Our first objective was to identify top hibernation-linked *cis-*elements and *cis-*regulatory programs for targeted functional studies. Previously, we showed that parallel accelerated regions in hibernators (pHibARs) accumulated in specific TADs (*28*). Therefore, using set of pHibARs and parallel hibernator deleted regions (pHibDELs) uncovered in our companion study, we tested for enrichments in TADs throughout the mouse genome compared to background conserved regions (CRs) (*27*). For comparison, we performed the same analysis for control homeothermic lineages that are the most closely related to the hibernators in the multiple genome alignment, identifying parallel homeotherm accelerated regions (pHomeoARs) and deleted regions (pHomeoDELs) (*27*). From 2153 TADs tested, we found 25 and 580 that are significantly enriched for pHibARs and pHibDELs, respectively (**Fig. 1A, blue**). In contrast, pHomeoARs and pHomeoDELs are significantly enriched in different TADs compared to hibernators (**Fig. 1A, red**). Therefore, our results identified TADs that disproportionately accumulated convergent genomic changes in hibernators at conserved *cis-*elements (**Table S1**).

**Fig. 1.**
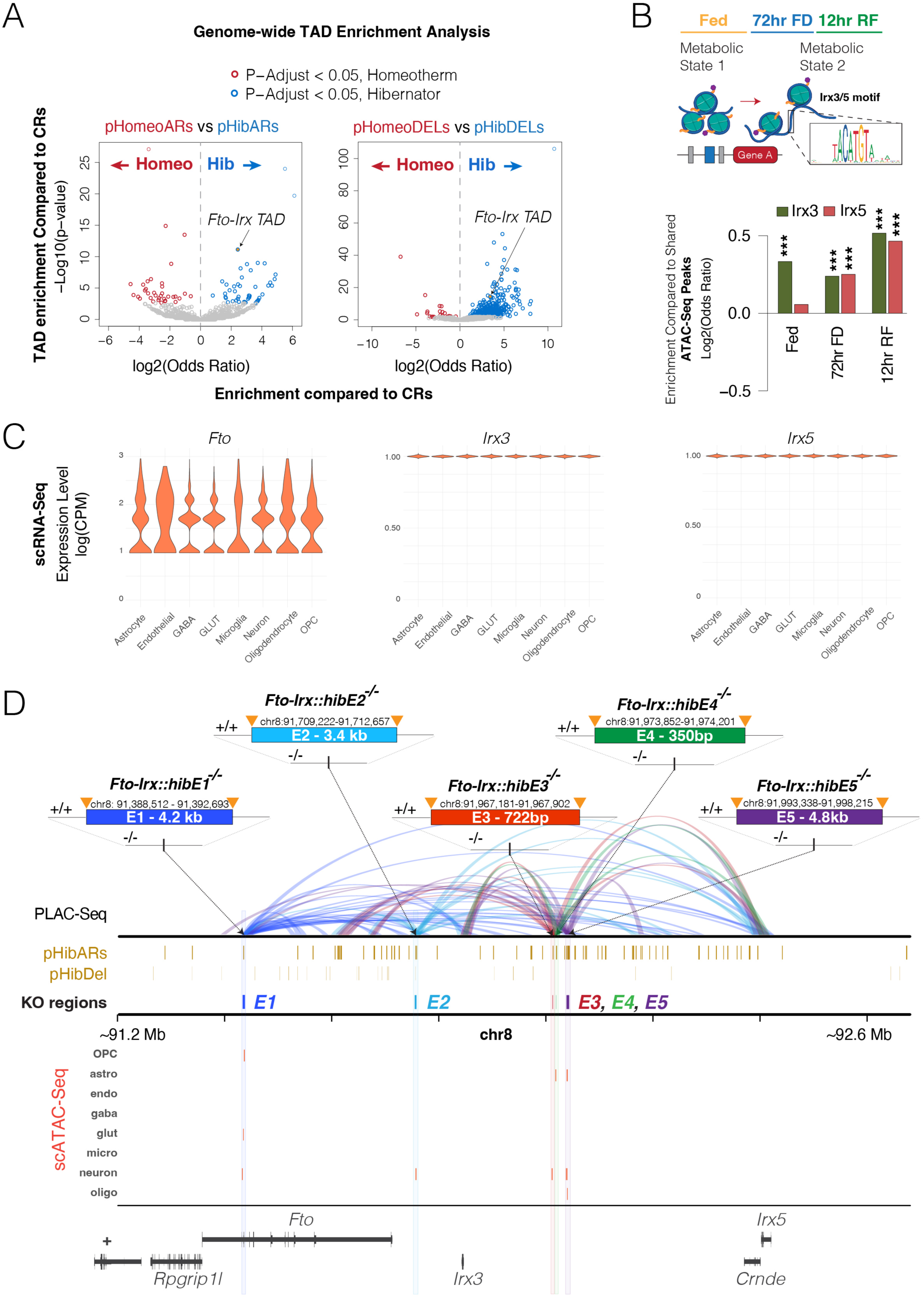
Convergent genomic changes in hibernators identify *Fto-Irx* TAD *cis-*elements for functional analysis in knockout mice. **(A)** Volcano plots show the odds ratio and p-values for TADs throughout the mouse genome to be enriched for hibernator (blue) and homeotherm (red) convergent genomic changes relative to CRs. The results in the left plot for pHibARs (blue) and pHomeoARs (red) are based on a chi-square test of a contingency table of AR and CR counts (rows) by hibernator and homeotherm data (columns). The same analysis for pHibDELs versus pHomeoDELs is shown in the right plot. Significantly enriched TADs for hibernator (blue circles) or homeotherm (red circles) convergent genomic changes are shown (q-value<0.05). Other TADs are indicated by grey circles. The Fto-Irx TAD is among the top enriched TADs for both pHibARs and pHibDELs. **(B)** Barplots show the odds ratio for *Irx3* and *Irx5* binding site motifs to be located in adult female mouse hypothalamus ATAC-Seq peak sites of open chromatin (FDR 5%) that are specific to Fed, 72 hr food deprived (FD), versus 12 hr refed (RF) conditions compared to peaks that are shared across all three metabolic conditions. The results show that Irx3 motifs are enriched in Fed, 72hr FD, and 12hr RF specific peak sites, while *Irx5* motifs are enriched in 72hr FD and 12hr RF specific peak sites. **(C)** The violin plots show adult hypothalamus scRNA-Seq data showing *Fto, Irx3,* and *Irx5* expression is present in all major cell-types of the hypothalamus. **(D)** The plot displays multi-omics data for the *Fto-Irx* TAD focused on 5 cis-elements showing convergent genomics changes in hibernators (*Fto-Irx::hibE1-5* cis-elements) that were selected for functional studies in knockout mice and the genomic size and coordinates for each *cis*-element. PLAC-Seq for H3K27ac shows the regulatory contacts made by each knocked out cis-element (E1, dark blue; E2, light blue; E3, red; E4, green; E5, purple) and reveals that each cis-element makes multiple long-distance contacts that affect *Fto, Irx3,* and *Irx5,* but not *Rpgrip1l*. Tracks (brown) are shown for the pHibARs and pHibDELs identified in the TAD and show the knockout targeted regions overlap with pHibARs (E1, E3, E4, E5) or pHibARs and pHibDELs (E2). The scATAC-Seq tracks show significant ATAC-Seq peaks in each targeted *cis-*element identified from single cell multi-omics profiling of fed, 72hr FD, and 12hr RF adult mouse hypothalamus (FDR < 5%). The open chromatin peaks for each targeted *cis-*element in oligodendrocyte precursor cells (OPCs), astrocytes (Astro), endothelial cells (endo), GABAergic neurons (gaba), Glutamatergic neurons (Glut), Microglia (micro), Other neurons (neuron), and Oligodendrocytes (Oligo). The results show that each *cis-*element shows significant chromatin accessibility in specific cell types of the hypothalamus.

Among the top pHibAR and pHibDEL enriched TADs, we identified the *Fto-Irx* TAD that includes *Rpgrip1l (RPGR-Interacting Protein 1-Like Protein)*, *Fto* (*Fat Mass & Obesity alpha-ketoglutarate dependent dioxygenase*), *Irx3* (*Iroquois 3 homebox transcription factor*), and *Irx5* (**Fig. 1A**). This finding using a new analytical strategy including Zoonomia data replicates our previous work (*28*), and unveils a top candidate TAD for functional studies of *cis-*elements impacted by pHibARs and pHibDELs. The *Fto-Irx* locus is the strongest genetic risk locus for human obesity (*34*), and affects age-related brain atrophy (*35*, *36*), feeding behaviors (*37–40*), and brain injury risk (*41*). Variants impacting a CRE in the first intron of *Fto* have been linked to obesity and affect the expression of neighboring genes, including *Irx3* and *Irx5,* through long range regulatory contacts (*30*, *34*). Our study uncovered 80 pHibARs and 37 pHibDELs that point to conserved *cis-*elements in the *Fto-Irx* region that could also regulate aspects of metabolism and behavior.

Convergent genomic changes in hibernators relate to gene expression programs for FFR responses in the hypothalamus (*27*). The regulatory functions of proteins encoded by *Fto*, a m6A demethylase, and *Irx3* and *Irx5* homeobox transcription factors, are compatible with potential roles in regulating FFR responses. To test this and determine if the *Fto-Irx* locus might be an important node in FFR response programs, we performed and analyzed ATAC-Seq profiles of chromatin accessibility sites in the hypothalamus for fed, 72-hr food deprived (FD), and 12-hr refed (RF) adult female mice. We then tested whether the DNA sequences for the open chromatin sites uniquely identified in Fed, 72-hr FD, or 12-hr RF conditions are significantly enriched for *Irx3* and/or *Irx5* binding site motifs compared to ATAC-Seq peaks shared across all three FFR conditions. We found that the *Irx3* motif is significantly enriched in open chromatin sites uniquely accessible in Fed, 72-hr FD, and 12-hr RF conditions, while the *Irx5* motif is significantly enriched in sites open in the 72-hr FD and 12-hr RF conditions (**Fig. 1B**). Thus, we found evidence for roles for transcription factors in the *Fto-Irx* TAD in broadly affecting FFR gene expression responses, providing further support for targeted functional studies of the hibernation-linked *cis-*elements in this genomic region.

To define individual *cis-*elements for targeted functional studies, we explored the *cis-*regulatory architecture of the *Fto-Irx* TAD in the mouse hypothalamus using H3K27ac+ PLAC-Seq, single cell (sc) multi-omics, and pHibAR and pHibDEL datasets (*27*). The scRNA-Seq data shows that *Fto, Irx3,* and *Irx5* are expressed in most major cell types in the hypothalamus (**Fig. 1C)**. Single cell ATAC-Seq data revealed pHibARs and/or pHibDELs that overlap noncoding sites showing chromatin accessibility in specific cell-types, and our PLAC-Seq data revealed genes with regulatory contacts from each *cis-*element. We identified 5 hibernation-linked *cis-*elements for targeted functional studies that demonstrate evidence for activity (open chromatin) in hypothalamic cells and regulatory contacts with multiple genes in the *Fto-Irx* TAD that suggest roles in coordinating *Fto, Irx3*, and *Irx5* expression (**Fig. 1D**). We named these *cis-*elements as follows: *Fto-Irx::hibE1, Fto-Irx::hibE2, Fto-Irx::hibE3, Fto-Irx::hibE4*, and *Fto-Irx::hibE5* (**Fig. 1D**). They display significant chromatin accessibility in hypothalamic OPCs, astrocytes, oligodendrocytes, glutamatergic neurons, and/or other neuron types in Fed, 72-hr FD, or 12-hr RF conditions (**Fig. 1D**). None show significant accessibility in microglia, GABA neurons, or endothelial cells. Each of the 5 *cis-*elements form multiple and distinct regulatory contacts across the *Fto-Irx* TAD (**Fig. 1D**). All 5 form contacts with *Irx3* and *Irx5* promoters and, with the exception of *Fto-Irx::hibE4*, form contacts with the *Fto* gene locus (**Fig. 1D**), suggesting different effects on *Fto, Irx3*, and *Irx5* expression in the hypothalamus and/or other tissues. None contact *Rpgrip1l*. We generated knockout mice for each *cis-*element to test functional roles in gene expression, metabolic, and behavioral control.

## *Fto-Irx::hibE*(*1–5*) Knockout Mice Exhibit Alterations to Hypothalamic Gene Expression and Molecular Responses to Fed and Fasting Conditions

We hypothesized that the *Fto-Irx::hibE cis-*elements regulate gene expression. To test this, we performed bulk RNA-Seq profiling to compare hypothalamic gene expression between *Fto-Irx::hibE* ^-/-^ and ^+/+^ adult female mice in fed and 72hr FD conditions. Each knockout mouse line showed significant gene expression changes compared to controls. We first focused on effects on the expression of *Fto, Irx3,* and *Irx5* genes and found that the ^-/-^ mice show significant changes to *Fto, Irx3,* and/or *Irx5* compared to ^+/+^ littermates in a fed and/or 72hr FD condition, with the exception of the *Fto-Irx::hibE1*^-/-^ mice, which did not show significant changes (**Fig. 2A-E**). *Fto-Irx::hibE2^-/-^* mice show decreased *Irx3* and *Irx5* expression in fed mice (**Fig. 2B**). *Fto-Irx::hibE3^-/-^* mice displayed increased *Fto* and decreased *Irx5* expression when fed, with no *Fto* change but reduced *Irx3* and *Irx5* expression in the 72-hour FD condition (**Fig. 2C**). In *Fto-Irx::hibE4^-/-^* mice, *Irx3* and *Irx5* expression reduced after 72 hours of FD (**Fig. 2D**), whereas *Fto-Irx::hibE5^-/-^* mice exhibited an increase in *Irx5* expression exclusively in the 72-hour FD condition (**Fig. 2E**). Significant changes to the expression of the neighboring *Rpgrip1l* and *Crnde* genes were not observed in any mice. These findings show that the 4 out of 5 *Fto-Irx::hibE cis-*elements affect *Fto*, *Irx3*, and/or *Irx5* expression in the adult hypothalamus in a metabolic state/food availability dependent manner.

**Fig. 2.**
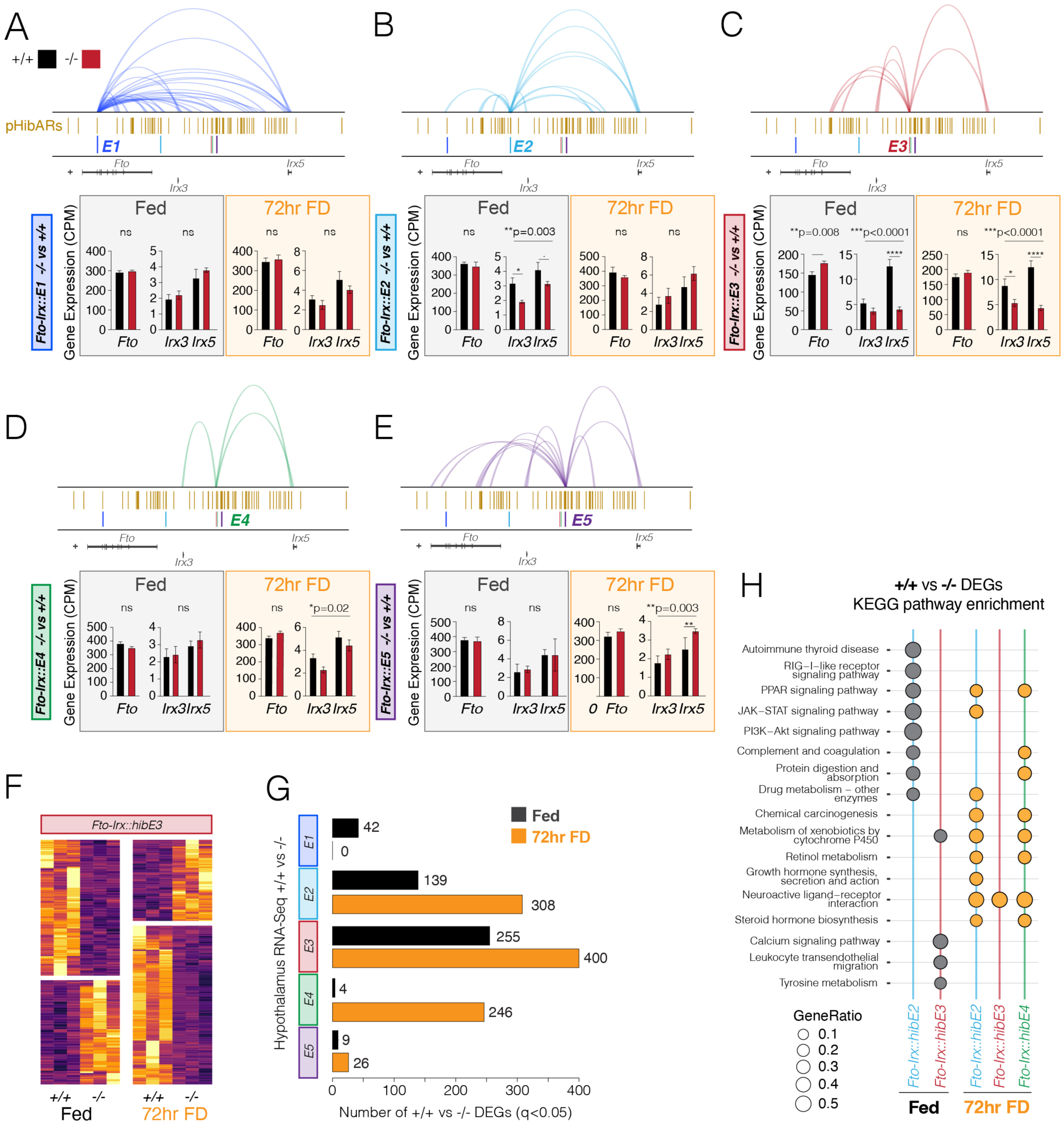
Gene expression alterations in the hypothalamus of *Fto-Irx::hibE cis-*element knockout mice under fed and 72-hr food deprived conditions. (**A-E**) The barplots show the RNA-Seq measured expression of *Fto, Irx3,* and *Irx5* in the hypothalamus in each of the *Fto-Irx::hibE1-5* knockout mouse lines (n=4, two-tailed test (*Fto*) or One-way ANOVA (*Irx3* and *Irx5*) with Tukey post-test). The H3K27ac+ PLAC-Seq detected regulatory contacts for each cis-element are shown above, along with tracks for pHibARs and the location of each knocked out *cis-*element (E). Data for mice in a fed (grey box) versus 72-hr FD condition (orange box) are shown. CPM, counts per million. The results show how each element affects *Fto, Irx3*, and *Irx5* expression in two different metabolic conditions. (**F**) The heatmap shows the expression of RNA-Seq detected differentially expressed genes (DEGs, FDR < 5%) in the hypothalamus for *Fto-Irx::hibE3*^-/-^ mice in Fed and 72 hr FD conditions relative to ^+/+^ littermates. (**G**) The barplots show the numbers of significant DEGs (RNA-Seq, FDR<5%) detected in the hypothalamus of ^-/-^ versus ^+/+^ mice for Fed and 72-hr FD conditions for all 5 different *Fto-Irx::hibE1-5 cis-*element knockout mice (E1-5). (**H**) The plot shows significant KEGG pathway enrichments from a gene set analysis of significant DEGs in *Fto-Irx::hibE2*^-/-^, *Fto-Irx::hibE3*^-/-^, and *Fto-Irx::hibE4*^-/-^ mice. Significant enrichments were not observed for DEGs in *Fto-Irx::hibE1^-/-^* or *Fto-Irx::hibE5^-/-^*mice.

Given their regulatory functions, changes to *Fto*, *Irx3*, and/or *Irx5* expression in our knockout mice could have downstream effects that impact wider gene expression programs. In support, we uncovered hundreds of differentially expressed genes (DEGs, FDR < 5%) in *Fto-Irx::hibE2*^-/-^, *Fto-Irx::hibE3*^-/-^, and *Fto-Irx::hibE4*^-/-^ mice relative to their respective ^+/+^ littermate controls (**Fig. 2F and G**). The numbers of significantly affected genes differed according to *cis-*element and metabolic state (**Fig. 2G**). Only modest effects were detected in *Fto-Irx::hibE1*^-/-^ and *Fto-Irx::hibE5*^-/-^ mice in either condition, suggesting that these elements do not strongly affect the adult hypothalamus (**Fig. 2G**). Gene set enrichment analysis of the sets of DEGs identified significant KEGG pathway enrichments that differed according to the *cis-*element and metabolic state of the animal (**Fig. 2H**). We observed significant enrichments in various signaling pathways, metabolic pathways, and hormonal pathways. Since the mice are germ-line knockouts, the gene expression changes could be caused by direct effects in the adult hypothalamus and/or by indirect effects from other tissues or development. Overall, our findings support functional CRE activity for conserved *cis-*elements impacted by pHibARs and/or pHibDELs and show that deleting a single hibernation-linked CRE in the *Fto-Irx* TAD can have effects on downstream gene expression.

## Hibernation-Linked CREs Modulate Metabolic Adaptations Relevant to Hibernation

Hibernation is an adaptation for surviving periods of food scarcity. Obligate hibernators seasonally regulate metabolism and behavior to gain fat (*8*), followed by extended periods of torpor involving suppressed metabolism, body temperature, and activity, and then emergence from torpor with refeeding and recovery (*5*, *6*). Mice do not hibernate but show brief fasting-inducible torpor bouts (*42*, *43*). We devised an assay of hibernation-relevant metabolic changes in mice to explore the phenotypes of our *Fto-Irx::hibE* CRE knockout lines. Adult female mice were implanted with small internal body temperature (Tbi) monitors and placed in CLAMS (comprehensive lab animal monitoring system) metabolic cages to measure calorimetric, activity, and feeding changes during a 7-day FFR paradigm that consists of (i) 48hrs in a fed baseline state (Pre-torpor phase), (ii) 48hrs of food deprivation (FD) + reduced ambient temperature (shift from 25°C to 18°C) for torpor induction (Torpor phase), and (iii) 3 days of refeeding (RF) at 25°C with ad libitum food (RF phase) (**Fig. 3A**). During the pre-torpor phase, mice show a Tb_i_ of 38°C and 36°C during the dark versus light cycles, respectively (**Fig. 3A, red pretorpor**). The Tb_i_ drops in response to FD + reduced ambient temperature, indicative of torpor bouts, reaching as low as ∼20°C (**Fig. 3A, blue torpor**). Within hours of RF, Tb_i_ returned to baseline pre-torpor levels (**Fig. 3A, orange refeeding**). With this FFR paradigm, we explored metabolic phenotypes in the different *Fto-Irx::hibE* knockout mice in pre-torpor, torpor, and refeeding conditions.

**Fig. 3.**
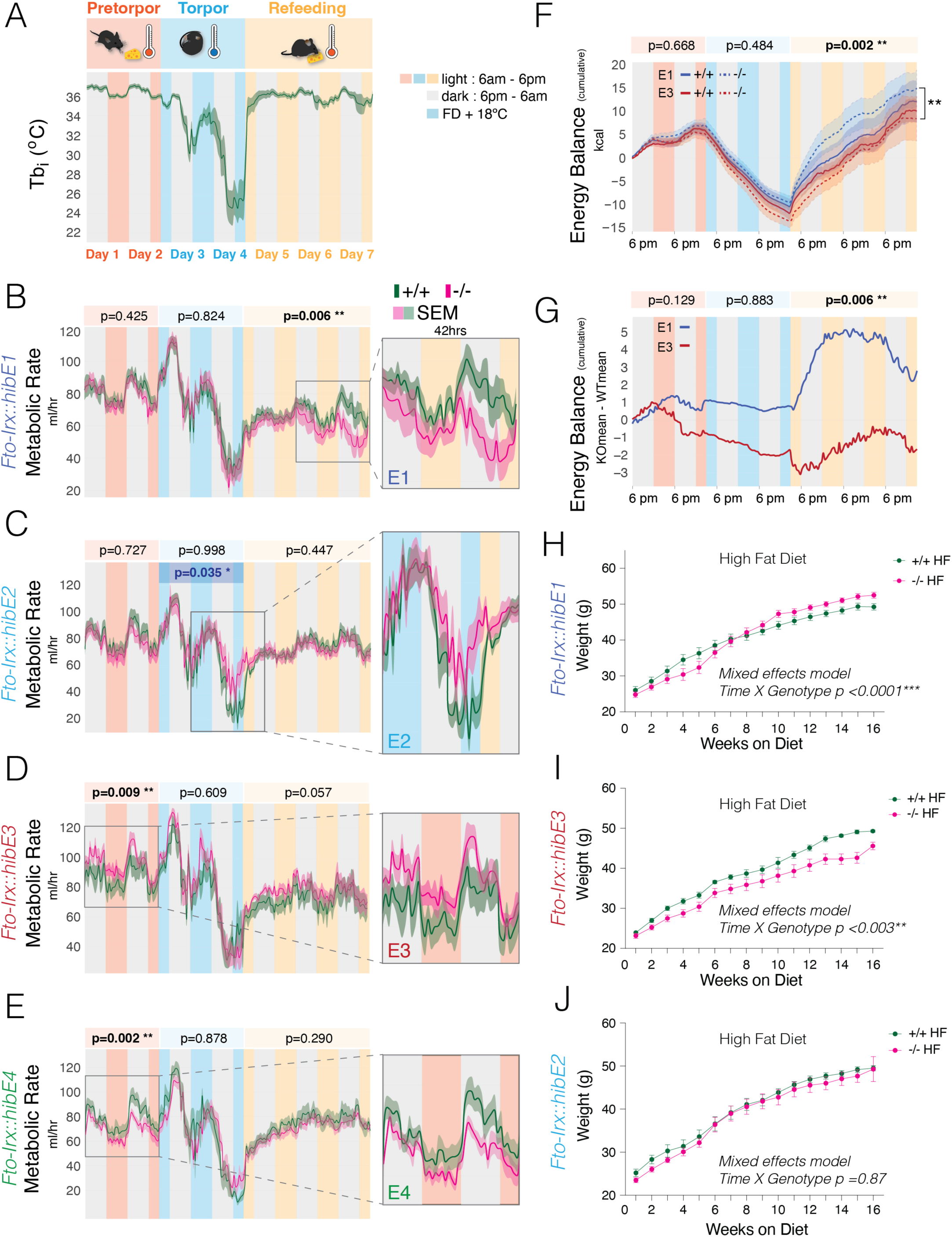
Metabolic and Thermoregulatory Response Profiles in *Fto-Irx::hibE cis-*Element Knockout Mice Across Nutritional States Mimicking Hibernation Cycles. (**A**) The plot shows internal body temperature (Tbi) in mice over 7-days while placed in CLAMS cages for a metabolic assay of processes pertinent to hibernation. The assay has a 48-hr pretorpor phase (red, ad libitum food + 25°C), 48-hr torpor phase (blue, FD + 18°C), and a 72-hr post-torpor refeeding phase (orange, ad libitum food + 25°C). Mice show circadian changes with elevated Tbi during active periods (dark cycle, grey bars) and depressed Tbi during inactive periods (light cycle, colored bars). During the torpor phase, Tbi drops, which is indicative of torpor, and then rapidly recovers during refeeding. Dark green line shows the mean, while the shaded area shows S.E.M. (**B-E**) The plots show the metabolic rate (MR, (O_2_ consumption ml/hr)) measured from ^-/-^ (pink) versus ^+/+^(green) adult female mice in CLAMS cages during the metabolic assay. Mean MR results (solid line) are shown and S.E.M. (shared area) for *Fto-Irx::hibE1*^-/-^ mice (n=11-14) (**B**), *Fto-Irx::hibE2*^-/-^ mice (n=7-8) (**C**), *Fto-Irx::hibE3*^-/-^ mice (n=7) (**D**), and *Fto-Irx::hibE4*^-/-^ mice (n=5-7) (**E**). Statistical significance in the pre-torpor, torpor, and refeeding phases was determined using a generalized linear regression analysis of the mean MR for each day and light-dark cycle (Day.LD) and tested for a main effect of genotype and an interaction effect between genotype and Day.LD. Variance due to batch effects from independent CLAMS cohorts was absorbed by a batch term in the model. P-values in black show the genotype effect (significant p-values in bold). P-values in blue show the genotype x Day.LD interaction effect (only significant p-values shown; C, torpor phase). *p<0.05, **p<0.01, ***p<0.001 (**F and G)** The plot in (F) shows the cumulative EB for *Fto-Irx::hibE1*^-/-^ mice (dotted blue line), *Fto-Irx::hibE1*^+/+^ mice (solid blue line), *Fto-Irx::hibE3*^-/-^ mice (dotted red line), and *Fto-Irx::hibE3*^+/+^ mice (solid red line). Dark lines show the mean, while the shaded area shows S.E.M. A generalized linear regression analysis of the mean EB for each day and Day.LD found a significant main effect of genotype (4 levels) in the RF phase, and a post-test showed that *Fto-Irx::hibE3*^-/-^ mice have a significantly reduced EB compared to *Fto-Irx::hibE1*^-/-^ mice (black line and asterisk on right side). This relative difference is made clear in (G) which shows a plot of the difference in the means of the cumulative EB for -/- minus +/+ mice and shows reduced EB in *Fto-Irx::hibE3*^-/-^ mice (red line) compared to *Fto-Irx::hibE1*^-/-^ mice (blue line). (**H-J**) The plots show the increase in body weight for -/- (pink) versus +/+ (green) adult male mice on an obesogenic high fat diet for *Fto-Irx::hibE1* (H; -/- n=8; +/+ n=10)*, Fto-Irx::hibE3* (I-/- n=10; +/+ n=5), and *Fto-Irx::hibE2* (J; -/- n=9; +/+ n=8). A significant genotype X time interaction was found for the *Fto-Irx::hibE1* and *Fto-Irx::hibE3* lines, with *Fto-Irx::hibE1^-/-^*mice gain more weight, while *Fto-Irx::hibE3^-/-^* gain less weight compared to ^+/+^ littermates (mixed model). A significant difference was not observed for *Fto-Irx::hibE2*.

Our study focused on 4 KO mouse lines, including those showing the strongest gene expression changes, *Fto-Irx::hibE2*^-/-^, *Fto-Irx::hibE3* ^-/-^, and *Fto-Irx::hibE4*^-/-^ mice, but also the *Fto-Irx::hibE1* ^-/-^ mice despite neglibible gene expression effects in the adult hypothalamus, because this *cis-*element is in the first intron of *Fto* near the region linked to human obesity (*30*, *44*). Adult female ^-/-^ mice were compared to littermate ^+/+^ controls and a regression analysis tested for a significant genotype effect and significant interaction effects between genotype and light-dark (LD) phase for each measure. Each CRE has significant and distinct phenotypic effects on metabolism *Fto-Irx::hibE1* ^-/-^ mice display significantly reduced metabolic rate (MR) during the RF phase (**Fig. 3B**); *Fto-Irx::hibE2*^-/-^ show increased MR during the end of the torpor phase that is detected from a significant interaction between genotype and LD (**Fig. 3C**); *Fto-Irx::hibE3*^-/-^ show significantly increased MR during the pretorpor phase(**Fig. 3D**); and *Fto-Irx::hibE4*^-/-^ show significantly decreased MR during the pretorpor phase (**Fig. 3E**). Our findings show that each CRE has significant and unique effects in modulating metabolism during pretorpor, torpor, or RF metabolic responses.

The full metabolic and activity profiles for each knockout mouse line during the assay further show that each CRE plays different roles in modulating metabolism (**Fig. S1-S4**). In addition to effects on MR, *Fto-Irx::hibE1^-/-^* mice are also characterized by significantly reduced Tbi and energy expenditure (EE) during the RF phase (**Fig. S1A and E**). *Fto-Irx::hibE2^-/-^* mice show significantly increased EE during the torpor phase (**Fig. S2E**). Further, we found significantly decreased locomotion (**Fig. S2B**) and energy balance (EB) (**Fig. S2I**) during the latter torpor phase, as well as a spike of increased in food intake during first hours of the RF phase (**Fig. S2G**). These effects involve a significant interaction between genotype and LD. Over the whole assay, cumulative food intake was significantly reduced in *Fto-Irx::hibE2^-/-^*mice. The effect is largely linked to a trend for reduced food intake during pre-torpor (**Fig. S2H**). Thus, while the *Fto-Irx::hibE1* CRE predominantly has effects during RF, the *Fto-Irx::hibE2* CRE has effects primarily during the deepest torpor stages and the emergence from torpor.

For the *Fto-Irx::hibE3^-/-^* mice, in addition to significantly increased MR, the full metabolic profiles also show significantly increased EE during pre-torpor (**Fig. S3H**), as well as increased cumulative EE over the whole assay (**Fig. S3F**). Significantly increased cumulative food intake during pre-torpor was also found (**Fig. S3H**), as well as reduced Tbi during RF (**Fig. S3A**). In contrast, *Fto-Irx::hibE4^-/-^*mice show the opposite phenotypic effects by exhibiting significantly decreased MR (**Fig. S4C**) and EE (**Fig. S4E**) during pre-torpor. These mice also show significantly decreased respiratory exchange ratio (RER) during the RF phase (**Fig. S4D)**. This indicates reduced CO2 production relative to O2 consumption, and decreased cumulative EE during torpor (**Fig. S4F)**, and decreased cumulative food intake and EB during RF (**Fig. S4H and J**). Thus, the *Fto-Irx::hibE3* and *Fto-Irx::hibE4* CREs are neighboring *cis-*elements that modulate pre-torpor, torpor, and RF metabolic phenotypes in opposing and different ways.

Prior to the assay, we found no significant body weight or composition differences between ^-/-^ and ^+/+^ mice for any of the four lines tested (**Fig. S5A-D**). This suggests that the significant metabolic phenotypes are not related to differences in body weight or composition. At the end of the assay, *Fto-Irx::hibE1^-/-^* mice showed a significantly reduced body weight, indicating weight loss and a failure to fully recover body weight after torpor (**Fig. S5A, post-assay body weight**). Significant body weight or composition changes were not detected in the other ^-/-^ mouse lines at the end of the assay, (**Fig. S5A-D**). Overall, each of the hibernation-linked CREs modulates metabolism differently.

Hibernators gain fat prior to torpor, pointing to changes to EB, feeding, and obesity control compared to homeotherms. EB measures the budget between calories ingested (kcal of food consumed) and calories expended (kcal of energy expenditure), and obese versus lean phenotypes are influenced by even small net imbalances between caloric intake and output over time. We observed that *Fto-Irx::hibE1^-/-^* (**Fig. S1J**) and *Fto-Irx::hibE3^-/-^* (**Fig. S3J**) mice show opposite trends in cumulative EB phenotypes over time. A regression analysis of the cumulative EB data for the two mouse lines confirmed that *Fto-Irx::hibE1^-/-^* mice have significantly elevated cumulative EB compared to *Fto-Irx::hibE3^-/-^* mice (**Fig. 3F and G**). Given this, we explored how these CREs affected weight gain responses to a high fat (HF) obesogenic diet in adult male mice over 16 weeks, contrasting the results with *Fto-Irx::hibE2^-/-^* mice that do not show trends for changes to cumulative EB (**Fig. S2J**). We found that *Fto-Irx::hibE1^-/-^*mice on a HF diet became significantly heavier than^+/+^ littermate controls (**Fig. 3H**), while *Fto-Irx::hibE3^-/-^* mice are significantly lighter than controls (**Fig. 3I**). In contrast, *Fto-Irx::hibE2^-/-^* mice were not significantly different from controls (**Fig. 3J**). The weight gain phenotypes agree with the cumulative EB trends for each ^-/-^ mouse line and demonstrate that each CRE has distinct effects on diet-induced weight gain.

Body composition analysis following the HF diet found that the *Fto-Irx::hibE1^-/-^*mice show a significant decrease in % lean body mass at the end of the HF diet compared to the beginning (**Fig. S6A**), while the *Fto-Irx::hibE3^-/-^* HF diet mice have significantly reduced % fat mass and increased % lean mass compared to ^+/+^ controls (**Fig. S6B**). No significant changes were observed in *Fto-Irx::hibE2^-/-^* mice (**Fig. S6C**). In summary, convergent genomic changes in hibernators pinpointed conserved CREs with functional roles in modulating pre-torpor, torpor, and refeeding metabolic phenotypes, as well as HF diet-induced weight gain.

## A Single Hibernation-Linked CRE Affects Foraging Behavior

Metabolism and foraging are intertwined in mammalian ecology and biology. Metabolism dictates the energetic needs of an organism, while foraging is an adaptive response and behavioral repertoire evolved to meet these metabolic demands (*45–47*). Metabolic phenotypes also strongly influence aging processes (*2*, *3*). Variants and genes in the *Fto-Irx* TAD have been linked to feeding behavior changes and age-related brain atrophy (*35–37*, *39*, *40*, *48*). Additionally, hibernators show behavioral changes compared to non-hibernators by seasonally foraging to gain weight and entering torpor during periods of food scarcity, rather than evolving adaptations for year-round foraging. Hibernators also show improved longevity and neuroprotection (*5*, *14*, *17*, *49*). Therefore, hibernation-linked CREs in the *Fto-Irx* TAD that influence metabolism could also influence foraging behaviors, as well as age-related foraging changes. To test this in an ethologically-relevant manner, we focused on *Fto-Irx::hibE3^-/-^* mice, because they showed the strongest gene expression and metabolic phenotypes, and used an assay and computational ethology approach that we previously developed to study the genetics of naturalistic foraging (*50–52*). Our foraging assay includes a foraging arena attached to a home cage (**Fig. 4A**). Mice are permitted free access to the arena for spontaneous foraging in two phases: (1) a 30-minute Exploration Phase where the animal explores a new environment and discovers a food patch (Pot 2), and (2) a 30-minute Foraging Phase 4hrs later in which the food is moved to a different patch (Pot 4) and buried in sand (**Fig. 4A**). The Exploration phase examines behavior in a novel environment, while the Foraging Phase examines behavior in a now familiar environment, where the mouse expresses perseverative memory responses of the former food patch (Pot2) (*52*). During this assay, mice perform many different round-trip foraging excursions from the home (**Fig. 4B**). We examine gross foraging behaviors, as well as more fine-grained behaviors uncovered from discrete round trip foraging excursions (**Fig. 4B**) (*50–52*). With this approach, we profiled foraging in male and female *Fto-Irx::hibE3^-/-^*and ^+/+^ mice for both young adult (2-3 month old) and aged (12-month old) cohorts.

**Fig. 4:**
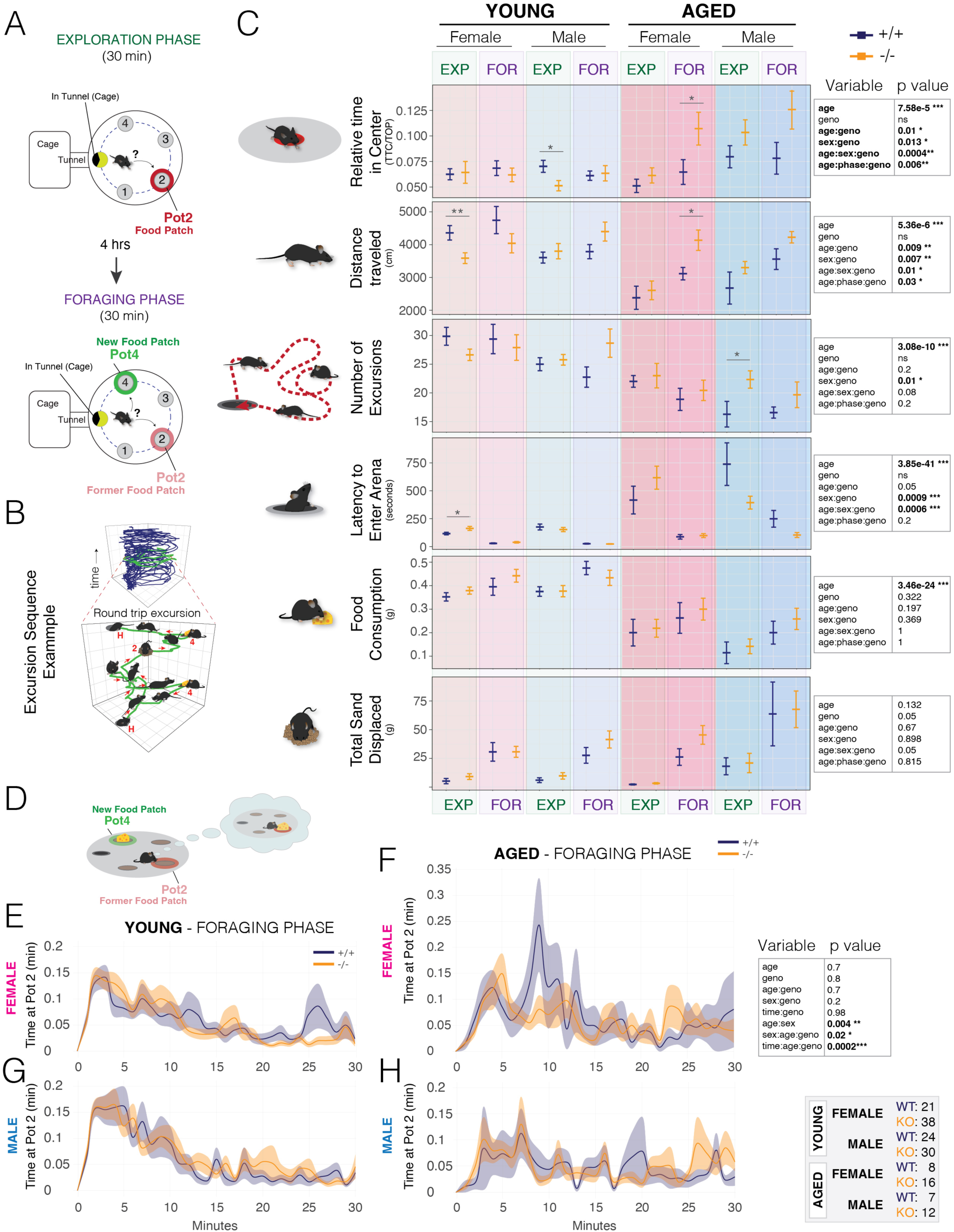
*Fto-Irx::hibE3* knockout mice exhibit significant changes to foraging behavior in a sex and age dependent manner. (**A and B**) The schematic shows the foraging assay design, including the home cage attached to the arena by a tunnel (A). The arena has 4 potential food patches and one (Pot2) has food in the 30-min naïve Exploration phase (red pot). In the 30-min Foraging phase 4 hours later the food is moved and buried in Pot4 (green pot), such that mice express memory response behaviors. Foraging behaviors are tracked and segmented into discrete round-trip excursions from the home and then decomposed by 52 measures of behavioral components (B). (**C**) The plots show data for 6 basic foraging behaviors expressed by 8-12 week old (young; n=21-38) and 12 month old (aged; n=7-16) males (blue) and females (pink) for *Fto-Irx::hibE3^-/-^* (orange) mice and ^+/+^ control littermates (blue). Multiple regression analysis revealed significant (bold type) main effects of age and/or sex on foraging measures, as well as significant interaction effects with genotype and phase (Exploration versus Foraging phase) (Multiple regression result tables are shown on the right). Post-tests identified specific contrasts with significant differences. Bars show mean ± SEM. Numbers of animals in each group are shown in the grey table on the bottom right. (**D-H**) The schematic depicts second-guessing perseverative behavior in the Foraging phase in which the mouse repeatedly investigates the former food patch learned in the Exploration phase (Pot2, red), rather than consuming readily available food in the new food patch (Pot4, green) (D and see Video S1). The plots show the time spent at Pot2 over the 30-min Foraging phase assay for young (E,G), aged (F,H), male (G,H), and female (E,F) ^-/-^ versus ^+/+^ mice. A table of the results of a multiple regression analysis using mixed models that test main effects of age, sex, genotype, and time, and interaction effects are shown on the right. Numbers of animals in each group are shown in the grey table to the right. Significant interaction effects between genotype, age, and sex are observed, as well as between genotype, age, and time. Aged female ^+/+^ mice show a large secondary wave of increased time at Pot2 (aka. second-guessing memory response) that is not observed in ^-/-^ aged females (F). Solid lines show the mean. Shaded area shows SEM. ***p<0.001; **p<0.01; *p<0.05.

We first examined effects on 6 gross foraging behaviors with a multiple regression analysis that tests for age, genotype, sex, and phase (Exploration versus Foraging) effects, as well as interaction effects. We found that all 6 behavioral show significant age effects, and 4 out of 6 measures show significant genotype (^-/-^ vs ^+/+^) effects that interact with age, sex, and/or phase (Exploration versus Foraging) (**Fig. 4C**). Therefore, loss of the *Fto-Irx::hibE3* CRE significantly influences foraging. Aged female and male ^-/-^ mice spend significantly more time in the exposed center during the Foraging phase (**Fig. 4C, Relative Time in Center**). Young female ^-/-^ mice travel a shorter distance during the Exploration phase, while aged female ^-/-^ mice travel an increased distance during the Foraging phase (**Fig. 4C, Distance Traveled**). Aged male ^-/-^ mice perform significantly more foraging excursions in the Exploration phase than controls (**Fig. 4C, Number of Excursions**). Young female ^-/-^ mice show a significantly increased latency to leave the home and forage during the Exploration phase (**Fig. 4C, Latency to Enter Arena**). Food consumption during foraging decreased significantly with age, but significant genotype effects were not observed (**Fig. 4C, Food consumption**). Sand displaced from digging was not significantly changed (**Fig. 4C, Total Sand Displaced**). Overall, this initial analysis of gross behavioral measures showed that loss of the *Fto-Irx::hibE3* CRE significantly influences foraging in a sex, phase, and age dependent manner. While significant, the phenotypic changes are complex. We therefore performed deeper analyses to better understand the affected behavioral components.

Our first deep analysis focused on memory response behaviors in the Foraging phase. Here, mice show repeated investigations of the former but now empty food patch (Pot2) (**Fig. 4D**, **Movie S1**). This “second-guessing” memory response is regulated by the synaptic plasticity gene, *Arc,* demonstrating mechanistic links to learning & memory (*52*). As previously described, we examined second-guessing responses in the mice by analyzing the cumulative time spent at the former food patch, Pot2, in 1-min time bins across the 30-minute Foraging phase (*52*). We found a significant interaction between genotype, sex, and age, indicating that *Fto-Irx::hibE3^-/-^* mice exhibit changes to second guessing in age and sex dependent manner. We also found a significant interaction between genotype, age, and time bin, which shows that the effects of the *Fto-Irx::hibE3* CRE depend on specific temporal epochs of the assay. Plots of the data show that aged female ^+/+^ mice show a pronounced peak in second-guessing between 5-12 minutes in the assay that is not displayed by the aged ^-/-^ females, nor by young adult females (**Fig. 4E and F**). Males also do not show this age-related behavioral change (**Fig. 4G and H**). Thus, age-related increases in a second-guessing memory-response behavior in females are absent from the knockouts, thereby defining a specific behavioral component affected by the *Fto-Irx::hibE3* CRE. We next investigated potential effects on discrete types of foraging excursions.

## *Fto-Irx::hibE3* Affects the Expression of Discrete Foraging Excursion Types Across the Lifespan

Foraging is a complex behavior, but genetic effects can be linked to finite behavioral sequences delineated from round trip excursions from the home that are described by 52 different measures of gait, velocity, movement trajectories, locations, and more (*50–52*). We found that the mice in our study expressed a total of 3,928 round trip foraging excursions (n=156 mice). Unsupervised hierarchical clustering and analysis of these with our DeepFeats algorithm identified 76 types of stereotypic, reproducible foraging excursions, which we call foraging “modules” (**Fig. 5A** and **Fig. S7A-F**). Forty-five foraging modules are expressed in both the Exploration and Foraging phases, while 13 and 18 are unique to the Exploration and Foraging phases, respectively (**Fig. 5A**). To confirm that excursions involving similar behaviors are classified to the same module, we visualized the full map of excursions and modules using PHATE (*53*) (**Fig. 5B, see full 3D map in Movie S2**). In this map, each dot is a single excursion embedded in space based on the 52 measures and excursions are colored by module assignments. Excursions classified to the same module are closely clustered (**Fig. 5B and Movie S2**). Thus, the modules capture related behaviors. Having defined and confirmed the foraging modules, we explored changes to module expression in *Fto-Irx::hibE3 ^-/-^* mice.

**Fig. 5.**
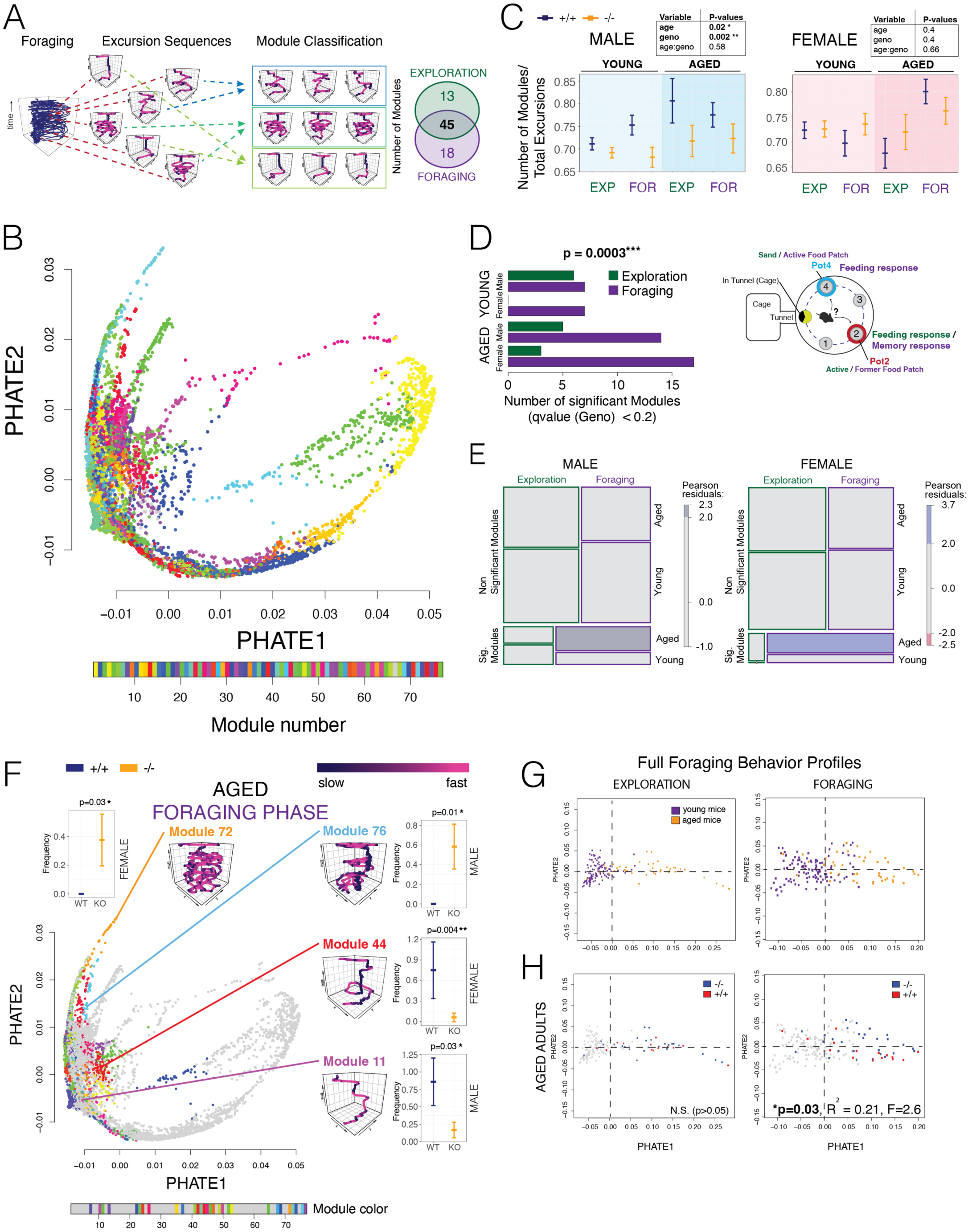
Differential expression of foraging modules in *Fto-Irx::hibE3^-/-^* mice and changes to the trajectory of behavioral aging primarily involves a memory-dependent foraging context. (**A** and **B**) The schematic overview depicts how different foraging excursions are grouped into subtypes stereotypic foraging excursions called modules using our unsupervised machine learning approach (A). The PHATE map shows 2-D embeddings of each excursion colored according to foraging module classification (B). The embeddings for each excursion are based on data from 52 different measures and the results show that excursions classified to the same module are closely clustered (see 3-D embeddings in Movie S2). The numbers of foraging modules uncovered from the young and aged, male and female, *Fto-Irx::hibE3* -/- and +/+ mice are shown in the Venn diagram for the Exploration and Foraging phases. (**C**) The plots show the number of different modules expressed by the mice normalized to the total number of excursions expressed, which is one measure of the diversity of different foraging strategies expressed. Multiple regression analysis revealed significant (bold type) main effects of age and genotype in males, as well as significant age*genotype interaction effects (Multiple regression results tables shown on upper right, n=15). Bars show mean ± SEM. (**D and E**) The barplot shows the number of significant differentially expressed modules between ^-/-^ and ^+/+^ mice in Exploration and Foraging phase for young and aged adults (q<0.2) (D). We observed that significantly more differentially expressed modules were found in the memory-dependent Foraging phase compared to the Exploration phase for aged males and females (linear model, phase*age interaction effect, p=0.0003). This age by phase dependence is shown for both sexes in mosaic plots for the counts of significant differentially expressed modules (E). (**F**) The PHATE map plots show the modules and excursions that are significantly differentially expressed in ^-/-^ versus ^+/+^ mice in aged adults (G) in the memory-dependent Foraging phase. The purple traces show foraging movement patterns representative of each affected example module shaded by velocity. Regression analysis revealed significant effects of genotype on the expression of these modules and the plots show the expression frequency of each foraging module in ^+/+^ versus ^-/-^ mice. Plots show the mean and error bars show SEM for the module expression frequency per mouse in *Fto-Irx::hibE3* WT (+/+, blue) and KO (-/-, orange) male or female mice (as indicated). (**G and H**) The plots show PHATE embedding of individual mice based on full foraging behavior profiles (basic measures + module expression) for the Exploration phase and Foraging phase data (G), where young adult mice (purple dots) and aged adult mice (orange dots) are shown, revealing that foraging profiles point to a trajectory of behavioral aging. PCA regression analysis of the full foraging behavior profiles for each mouse found that foraging behaviors are significant predictors of genotype (-/- (blue dots) versus +/+ (red dots)) in the aged adults in the memory-dependent Foraging phase (H, Foraging), but not the Exploration phase (H, Exploration). Young adults are shown by grey dots (H). See table in Figure 4 for number of mice per group. ***p<0.001, **p<0.01, *p<0.05.

We found that *Fto-Irx::hibE3 ^-/-^* male mice exhibit significantly fewer modules relative to the total number of excursions performed than ^+/+^ littermates (**Fig. 5C**). No significant difference was observed in females. This shows a sex difference and that loss of the *Fto-Irx::hibE3* CRE reduces the diversity of foraging modules expressed by male mice across tested ages, revealing a reduced repertoire. To examine effects on specific modules, we tallied the expression frequency of each module for each mouse and performed module-wise testing for a genotype effect (^-/-^ versus ^+/+^). Following multiple testing correction, we identified significant differentially expressed foraging modules between ^-/-^ and ^+/+^ mice for males and females in the Exploration and Foraging phases (**Fig. 5D**). Aged ^-/-^ males and females show more differentially expressed modules compared to^+/+^ controls than young adults, and that the effect is largely in the Foraging phase that tests perseverative memory responses (**Fig. 5D**). Indeed, a regression analysis of the numbers of significant modules found a significant interaction effect between genotype, age, and phase (Exploration versus Foraging) (**Fig. 5D**). The phase and age dependent effects on the numbers of significant modules are apparent in mosaic plots for males and females (**Fig. 5E**). Thus, loss of *Fto-Irx::hibE3* affects foraging modules in both sexes, with the most affected modules occuring in the memory-dependent Foraging phase in aged mice.

We visualized the differentially expressed modules in young (**Fig. S8 and Movie S3**) and aged adults (**Fig. 5F and Movie S4**) for the Foraging phase in PHATE maps. The results show that diverse module types across the map are affected, rather than a set of similar modules grouped in one region of the map. In aged mice, loss of *Fto-Irx::hibE3* significantly affected the expression of a range of different module types, including modules for long, complex exploratory behaviors (**Fig. 5F**, Module 72 and 76), goal-directed modules in which mice focus on the new food patch (Pot4) (**Fig. 5F**, Module 44), and brief darting behaviors involving briefly entering and exiting the arena from the home (**Fig. 5F**, Module 11). On the other hand, ∼43.4% modules (33 out of 76) were not significantly affected at any age or phase (**Fig. 5F and Movie S3,** grey dots). Therefore, the *Fto-Irx::hibE3* CRE significantly modulates the expression of a subset of foraging modules across the lifespan.

In a final analysis to test effects on the trajectory of behavioral aging, we aggregated the 6 gross foraging measures with the module expression data to create foraging profiles for each mouse. PHATE projections of the data show that the young and aged mice are differentiated in the Exploration and Foraging phases, revealing a trajectory of age-related changes in each context (**Fig. 5G**). PCA logistic regression analysis found that the behavior profiles significantly predict the -/- versus +/+ genotypes of aged mice in Foraging phase (**Fig. 5H**, p=0.03), and that PC4 (p=0.0086) and PC5 (p=0.049) are the significant predictors. The model explains about 21% of the variance; other unknown variables are needed to fully explain all the variance between two genotypes. Therefore, a singular hibernation-linked *cis-*element that modulates metabolism also significantly shapes naturalistic foraging and age-related changes, demonstrating that these processes are intertwined at the level of individual CREs.

## DISCUSSION

Our study illuminates genetic determinants of metabolic control. We show functional effects from deletions of single hibernation-linked *cis-*elements on gene expression, metabolism, and foraging in mice. The *Fto-Irx* TAD disproportionately accumulated convergent genomic changes in hibernators, defining hibernation-linked *cis-* elements that overlap accessible chromatin sites in the hypothalamus and form regulatory contacts with *Fto, Irx3,* and *Irx5*. Knockout mice for 5 different *Fto-Irx::hibE cis-*elements revealed that each modulates the expression of these 3 genes differently with indirect effects on the expression of hundreds of genes in a metabolic-state dependent manner. Assays of pre-torpor, torpor, and refeeding metabolic phenotypes showed that each targeted *cis-*element modulates different aspects of metabolism, with significant and unique effects on MR, Tbi, EE, EB, and locomotion control for hibernation-relevant states. The *Fto-Irx::hibE1^-/-^* and *Fto-Irx::hibE3^-/-^ cis-*elements affected diet-induced weight gain, which is key for hibernators to gain fat before torpor. By studying naturalistic foraging in young and aged adult *Fto-Irx::hibE3^-/-^* mice, we found that a single hibernation-linked CRE modulates specific aspects of foraging in a sex dependent manner, and affects the trajectory of age-related changes to foraging. Thus, metabolism and foraging are biologically intertwined at the level of hibernation-linked CREs. Conserved *cis-*elements discovered from convergent genomic changes in hibernators identify functional CREs for metabolic and behavioral adaptations involved in surviving challenging environmental conditions.

Mammalian genomes are organized into hierarchically structured TADs that contain DNA loops that control the regulatory specificity of enhancers and their interactions with gene promoters within the TAD (*54*, *55*). TADs are largely conserved across mammals, while the DNA loops within TADs are faster evolving, cell-type specific, and regulatory changes contribute to species differences in gene expression (*56–59*). Building on our previous work (*28*), we show that a subset of TADs disproportionately accumulated convergent genomic changes in hibernators (pHibARs and pHibDELs) pointing to TADs predicted to show changes to *cis*-regulatory architecture important for hibernation-linked traits. Our companion study found that most accelerated regions (and deleted regions) involve loss-of-function effects in which *cis-*elements conserved and showing activity in non-hibernators experience changes due to relaxed purifying selection in a hibernator (*27*). Thus, we can infer that TADs enriched for pHibARs and pHibDELs harbor *cis-*elements that are no longer essential in hibernators, and their elimination potentially enables new dynamics in metabolic, Tb_i_, and behavioral control. The *Fto-Irx* TAD contains many different CREs (*29*). We found 80 pHibARs and 37 pHibDELs in the *Fto-Irx* TAD that point to *cis-*elements conserved in homeotherms and changed in hibernators. Each *cis-*element could function in different tissues, cell-types, physiological contexts, and developmental stages. Our targeted functional analysis of 5 *cis-*elements found effects on the expression of multiple neighboring genes, including *Fto, Irx3*, and/or *Irx5,* in the hypothalamus in a metabolic state dependent manner. This uncovered roles in coordinating the expression of multiple genes suggesting they are hubs of regulatory control in the TAD. The *FTO* variant linked to human obesity lies in a CRE that also affects the expression of multiple neighboring genes in the TAD through long range regulatory contacts (*30*, *60*), highlighting the fact that individual CREs can affect broader genetic programs. Our findings further support this by showing that *Fto-Irx::hibE cis-*element knockout mice display significant changes to the expression of hundreds of genes in the adult hypothalamus in specific pathways. Thus, individual hibernation-linked CREs can be key genetic elements orchestrating entire genetic programs.

Hibernating species evolved extreme metabolic and behavioral adaptations for gaining fat and entering long periods of torpor with suppressed Tbi, metabolic rate, and energy utilization (*5*). Our knockout mouse studies show that different hibernation-linked *cis-*elements modulate different aspects of metabolism during pre-torpor, torpor, and post-torpor RF periods, and have distinct effects on weight gain responses to a HF diet. *Fto-Irx::hibE1^-/-^* mice show decreased MR during post-torpor RF, *Fto-Irx::hibE2^-/-^* mice show increased MR during torpor, *Fto-Irx::hibE3^-/-^* show increased MR during pre-torpor, while *Fto-Irx::hibE4^-/-^* mice show the opposite with decreased MR during pre-torpor. These phenotypes, as well as other significant metabolic and activity phenotypes in our assay of hibernation-relevant traits, reveal that each CRE plays different roles in modulating aspects of metabolism. The same conclusion was found for HF diet-induced obesity, where *Fto-Irx::hibE1^-/-^* mice became significantly heavier than controls, *Fto-Irx::hibE3^-/-^* mice were significantly lighter, and *Fto-Irx::hibE2^-/-^* mice were not significantly affected. Therefore, in a manner analogous to how CREs can shape aspects of animal morphology (*20*, *21*), our findings show that hibernation-linked CREs can have significant phenotypic effects that shape varied aspects of metabolism.

Metabolism, foraging, and lifespan are biologically and ecologically interconnected. Hibernating species have honed their foraging strategies to capitalize on high-caloric food sources in preparation for periods of torpor, contrasting with non-hibernators that exhibit diverse search strategies, dietary versatility, and adaptive foraging behaviors suited to fluctuating environmental conditions (*5*, *10*). In addition to behavior changes, hibernators exhibit enhanced longevity, neuroprotection, and tauopathy reversal (*5*, *14*, *17*, *49*, *61*). The *Fto/FTO* locus affects reward learning (*39*, *62*), locomotor responses (*40*), impulsivity (*37*), changes to food intake and preference (*63*), as well as age-related brain atrophy (*35*). Thus, we tested whether hibernation-linked CREs in the *Fto-Irx* TAD not only affect metabolism, but also show roles in ethologically relevant foraging phenotypes throughout life. Using a naturalistic foraging assay and computational methods that decompose foraging into components that can be tested for links to genetic factors, we studied young adult and aged *Fto-Irx::hibE3^-/-^* mice. Significant effects were uncovered at the level of gross measures, like total distance traveled, and at the level of more fine-grained behavioral components, including changes to “second-guessing” memory responses to a formerly learned food patch (*52*), and the expression of a subset of foraging modules that related to types of excursions from the home. The largest effects were uncovered in aged mice in a foraging context that involves memory-responses. Our results demonstrate that metabolism and foraging can be modulated by individual hibernation-linked CREs. Collectively, these studies offer insight into the genetics of metabolism and foraging.

## Acknowledgments

Thank you to Drs. Jared Rutter, Scott Summers, and all members of the Gregg lab for commenting on the manuscript. Knockout mice were designed and generation by the University of Utah Mutation Generation & Detection Core and Transgenic Mouse Core Facilities. The CLAMS metabolic phenotyping experiments were performed with the University of Utah Metabolic Phenotyping Core Facility. Bulk and single cell genomics experiments and sequencing were performed with the University of Utah High Throughput Genomics Core. Squirrel tissue was obtained from Dr. Dana Merriman (Squirrel Colony at the University of Wisconsin Oshkosh). Thank you to Tyler C. Leydsman and Bennett Cottle for technical and mouse colony support.

## Author contributions

Conceptualization: SS, EF, CG

Methodology: SS, CSH, EF, JE, CD, AC, AT, CG

Investigation: SS, CSH, EF, JE, JGM, ACR, TCL

Visualization: EF, CSH, SS, CG

Funding acquisition: CG

Project administration: CG

Supervision: CG

Writing – original draft: CG

Writing – review & editing: CSH, SS, EF, AC, AT, JGM, ACR, CG

## Data and materials availability

All data are available in the main text or the supplementary materials, and all genomics files are deposited in the NIH Short Read Archive repository.

## Supplementary Materials

Materials and Methods

Figs. S1 to S9

Movies S1 to S4

