## Supplementary Materials for "Conserved Noncoding Cis-Elements Associated with Hibernation Modulate Metabolic and Behavioral Adaptations in Mice"

**The PDF file includes:**

Materials and Methods
Figs. S1 to S8
Table S1 and S2
References

**Other Supplementary Materials for this manuscript include the following:**

Movies S1 to S4

**Materials and Methods**

### EXPERIMENTAL MODEL AND SUBJECT DETAILS

*Mice*

**Housing and Husbandry**

All experiments were conducted in compliance with protocols approved by the University of Utah institutional animal care and use committee (IACUC). FTO-pHibAR KO mice (see below) were bred and housed on ventilated racks at the University of Utah Comparative Medicine Center on a 12hr light cycle, 6am on and 6pm off, with an ambient temperature range of 20-25C. All mice were given water and food (Harlan-Teklad 2920X soy protein-free) ad libitum, with the exception of a single overnight fast for mice tested for foraging behavior (see METHOD DETAILS, Behavior, foraging), fasting periods for genomic and torpor experiments. Adult breeders (6weeks to 1year of age) were paired continuously, and pups were weaned at postnatal day 21 (P21) and cohoused with up to five same-sex littermates or similar aged same-sex mice of the same line; mice were never singly housed. As needed, ear punches were taken at P17-P21 for both genotyping biopsy samples and mouse identification purposes. Before dissections of brain tissue, mice were put to sleep with isoflurane gas anesthesia and decapitated.

**C57Bl/6J Mice**

C57Bl/6J mice for genomics experiments were commercially acquired from Jackson Labs (000664).

**pHibAR Knockout Mice**

CRISPR Cas9 reagents were designed and generated by the University of Utah Mutation Generation and Detection Core (see Table S1 for N20 target sequences). Off-targets were screened using CasOT (PMID: 24389662). For each pHibAR line a pair of sgRNAs, targeting just 5' and 3' of the target enhancer, were complexed with Cas9 protein (IDT) to form ribonucleoprotein (RNP) complexes that were electroporated into single cell mouse embryos by the University of Transgenic & Gene Targeting Mouse Core Facility (2x150ng sgRNA;100ng Cas9 protein). F0 founder mice were backcrossed into the C57Bl/6J background strain for five generations to reduce propagation of any potential unknown off-target CRISPR mutations, then subsequently maintained by breeding non-sibling heterozygous pairs to produce heterozygous, homozygous, and wild-type mice.

**Genotyping**

Ear punches taken at P7-P21 were lysed in "HotSHOT" lysis solution (Truett, G.E., et al. 2000) on a 95°C heat block for 30 minutes. Lysates were then pH neutralized with an equal volume of HotSHOT neutralizing solution. Two µL of lysates were then added to make 20µL PCR reactions with *Taq* DNA Polymerase (New England Biolabs, #M0495) and 2µM primer mixes (Table S2). Primers flanking the target enhancer deletion regions were used to detect the shorter mutant products and "WT" primers that bind within the target region were used to detect the WT allele to determine zygosity.

**Table S1: CRISPR-Cas9 sgRNA N20 sequences for targeting Fto-Irx enhancer regions. Each sgRNA were electroporated as a pair to induce fragment excision.**

| Enhancer | 5' N20 Sequence | 3' N20 Seqeunce | GRCm39 coordinates |
|---|---|---|---|
| Fto-Irx::hibE1 | CACCTATAAGGCAGCCGTGT | AAGTATCCGTATGTGGCGAA | 8:92114859-92119641 |
| Fto-Irx::hibE2 | CCGTCCTCAAGGCTGGGCCG | CCGGCACCTAGCGAGATGAG | 8:92435624 - 92439451 |
| Fto-Irx::hibE3 | TCCCTGGCCGGCGTGGAGAC | TTCCGTTTAGCTCGGAAGCA | 8:92693703 - 92694708 |
| Fto-Irx::hibE4 | ACTTAATCGGCCGGCTAAGT | TGTTAATGGCGCTGATGATG | 8:92700462-92700882 |
| Fto-Irx::hibE5 | AAGCAGAGCTAGCGAGCCTG | TATAGTAGGCTTGAGCCCAC | 8:92719751 - 92724937 |

**Table S2: Fto-Irx genotyping primer sequences with expected mutant and WT sizes in bps.**

| | **Flanking Forward** | **Flanking Reverse** | **WT Primer** | **Mutant** | **WT** |
|---|---|---|---|---|---|
| Fto-Irx::hibE1 | taggcaaggctgtcgtcgag | ccaaaacccccagccaacag | agccaccatgtcaccaacca | 828 | 478 |
| Fto-Irx::hibE2 | tccaaacagggatgcagcca | gccaggctgatgaaagtggc | gtcagtccctctgctgcctt | 545 | 287 |
| Fto-Irx::hibE3 | cccacctcattggccaggaa | gatcgccatcacctttctctct | ccgaccgttcactccgagta | 462 | 326 |
| Fto-Irx::hibE4 | agcaacctcagagctccacc | gtctctgtacgcggtgactgt | acaaaatcagcagcctggcc | 276 | 482 |
| Fto-Irx::hibE5 | tctgcaaccacagacagcac | gtgggacctctgtgctacct | gttctgagcgacctcacagc | 518 | 643 |

### METHOD DETAILS

Bulk Tissue Genomics

#### RNA-seq

Hypothalamus was dissected from adult female mice midday and RNA was isolated using the RNeasy Lipid Tissue kit (Qiagen 74804), treated with DNAse (Qiagen 79254). For food deprived mice, mice were moved into a clean cage and food was removed for 24-hrs, 72-hrs, or

72-hrs with reduced ambient temperature (18°C) prior to harvesting, with similar circadian timing to fed mice (midday). Refed mice were exposed to 72-hrs FD and then provided ad libitum access to mouse chow for 1hr (1hr RF), 12hrs (12hr RF), or 1 week (1 week RF). RNA-seq was performed by the Huntsman Cancer Institute High-Throughput Genomics Core at the University of Utah, using NEBNext Ultra II Directional RNA Library Prep with rRNA Depletion Kit (human, mouse, rat) library prep and NovaSeq S4 Reagent Kit v1.5.

**ATAC-seq**
At eight weeks of age, we dissected, and flash froze female mouse brain hypothalamic tissue for Fed, 72-hr FD, 12-hr RF, and 1-week RF (n=8). Samples were prepared for ATAC-Seq profiling of open chromatin using a commercial ATAC-Seq Kit (Active Motif, #53150).

**PLAC-seq**
8-10 week old adult female C57Bl6/J x CasteEiJ F1 hybrid mouse hypothalamus was dissected and flash-frozen (n=10). A ChIP-grade antibody for H3K27ac (AbFlex Histone H3K27ac rAb, Active Motif, cat #91193) was used. Samples were prepared for PLAC-Seq profiling of regulatory contacts using a commercial kit (Arima Hi-C kit+, Arima Genomics). Libraries were prepared for sequencing by the University of Utah High Throughput Genomics core facility using the NEBNext ChIP-Seq Library Prep Reagent Set for Illumina (E6200). Sequencing was performed using an Illumina NovaSeq 6000 with NovaSeq S1 Reagent Kit v1.5_150x150 bp paired end sequencing.

**H3K27ac ChIP-Seq**
H3K27ac 8-10 week old adult female mouse or adult female 13-lined ground squirrel hypothalamus was dissected and flash-frozen. A ChIP-grade antibody for H3K27ac (AbFlex Histone H3K27ac rAb, Active Motif, cat #91193) was used for chromatin immunoprecipitation. ChIP was performed using the ChIP-IT High Sensitivity kit (Active Motif, Cat# 53040). Libraries were prepared for sequencing by the University of Utah High Throughput Genomics core facility using the NEBNext ChIP-Seq Library Prep Reagent Set for Illumina (E6200). Sequencing was performed using an Illumina NovaSeq 6000 with NovaSeq S1 Reagent Kit v1.5_150x150 bp paired end sequencing.

Single cell Multi-Omics

**Hypothalamus single nucleus RNA-Seq + ATAC-Seq**
At eight weeks of age, we collected female mouse brain hypothalamic tissue for Fed (n=2), 72hr FD (n=3), and 12hr RF(n=3) conditions. We flashed freeze the tissue in liquid nitrogen and stored in -80 °C freezer. We follow the 10X GENOMICS nuclei isolation protocol specific for single cell multiome ATAC + Gene Expression (CG000505, PN-1000493, PN-1000494). In brief, tissue was submerged in 200 uL of Lysis Buffer and dissociated with pestle and incubate in ice for 10 minutes. Passing through a column, we centrifuge the dissociated tissue at 16,000 rcf for 20 seconds at 4 °C. Then, we discarded the column, vortexed the flow and centrifuge for additional 3 minutes at 500 rcf and 4 °C. We removed the supernatant, resuspended the pellet in 500 uL Debris Removal Buffer, and centrifuged at 700 rcf for 10 minutes at 4 °C. Finally, we discarded the supernatant, resuspended the nuclei pellet in 30 to 50 uL of Resuspension Buffer

and proceed to calculate nuclei concentration, nuclei quality assessment, library preparation, and sequencing. We prepared all buffer following the vendor's protocol.

The CellRanger - arc version 2.0.2 pipeline (10X Genomics) was used to process the scRNA-seq and scATAC-seq FASTQC files. The cellranger counts command was used to align the raw FASTQ files to the mm10mouse reference genome. For gene expression sequencing, filtered count matrices were analyzed using the R packager Seurat (V5.0.0) [1]. We emphasized in the designed for the integration of single-cell RNA sequencing (scRNA-Seq) datasets, specifically targeting the correction of batch effects in all replicates. Samples were merged into a single Seurat object to perform consistent filtering and quality-control (QC). Cells with low features or unique molecular identifier (UMI) counts or high mitochondrial read percentage were discriminated. In addition, to remove doublets, cells with abnormally high UMI counts were removed. Finally, any cells expressing mutually exclusive markers were removed. At the end, 61,400 cells passed filtering and QC processing. Data normalization was done using SCTransform R package, and dimensionality reduction techniques, including PCA and t-SNE, were applied. The data integration to correct batch involved the identification of integration features, anchors, and subsequent data integration using Canonical Correlation Analysis (CCA). The dimensional reduction t-SNE is applied uncovering 42 different hypothalamic cell subtype clusters.

Using the conserved markers function in Seurat and Celldex (v1.13) R packages, we identified eight different cell types: Astrocytes, Endothelial, GABA neurons, Glutaminergic neurons, Microglia, Other Neurons, Oligodendrocytes, and Oligodendrocyte Precursor Cells (OPC) and assigned their identity to the corresponded cluster. Consequently, we performed gene expression analysis using the conserved marker function of the Seurat package at between the different metabolic categories (FFR) at the cell level and cluster level.

<u>Metabolic Phenotyping</u>

**Body Composition and Weight**

For torpor experiments, body weight and composition were measured (NMR, Bruker Minispec LF50) prior to logger implantation and after logger removal post-CLAMS experiment. Body weight was also measured before mice were placed into CLAMS.

**Surgical Implantation of Body Temperature Logger**

At 2.5-5 month of age and body mass range of 18-25g, genotypically-balanced cohorts of pHibAR KO and WT female mice were surgically implanted with a temperature data logger (Star-Oddi, DST nano-T) programmed to record at 10-minute intervals. The mice were anesthetized with an isoflurane vaporizer, administered with lidocaine (0.003mg/g) and carprofen (0.005mg/g), and the logger was subcutaneously implanted through a 1cm mid-scapular incision and closed with surgical wound clips. Mice were administered with analgesics for two days post-op and singly housed for at least two weeks to recover, with food and water available *ad libitum*.

**Torpor Induction and Metabolic Monitoring**

The female pHibAR KO and WT mice with body temp implants were singly housed in metabolic CLAMS cages (Comprehensive Lab Animal Monitoring System, Columbus Instruments; University of Utah Metabolic Phenotyping Core) for total one week. Mice were allowed to acclimate to the cage, with regular food, at 25C for the first two days. On day 3 at 12pm, food was removed and the ambient temperature was dropped to 18C to induce torpor behavior. After 48hr, mice were refed and warmed back to 25C and monitored for another 72hr. At the end of the experiment, mice were euthanized for retrieval of the body temperature loggers and body weight and composition were again measured.

**Obesogenic Diet-Induced Phenotypic Screening in FTO-Modified Mice**

Young adult male (6-8 week old) *Fto-Irx::hibE* knockout mice and wild-type littermates were place on a high fat diet and weighed weekly for 16 weeks. Their body compositions at the beginning and end of the assay were measured, including fat, lean, and fluid weights, using a Bruker LF50 Nuclear Magnetic Resonance (NMR) system.

Dietary Intervention: The mice were randomly assigned to one of two dietary regimens provided by RESEARCH DIETS Inc.: a high-fat diet (60% kcal from fat; product number D12492) or a control diet comprising 10% kcal from fat (product number D12450J). Diets were administered ad libitum over a period of 22 weeks.

Data Analysis: The collected data on body weight and composition changes were analyzed using GraphPad Prism software using a two-tailed t-test or mixed model to determine the impact of genotype on body weight or composition.

Foraging Behavior Phenotyping

Foraging behavior testing was performed as detailed previously (*1*). In brief, in preparation for the foraging assay, mice were first habituated with sand (Jurassic play sand, Jurassic Sand) and seeds (Whole millet, Living Whole Foods) for two days in their home cage. On day one, seeds are spread on top of sand in the bottom of a Petri dish and the dish is placed on the bedding in the home cage for the mice to explore. On day two, seeds are covered with sand in the bottom of the Petri dish in the home cage for the mice to dig in and explore. To motivate animals to feed, mice were food deprived prior to testing to achieve 8%–10% weight loss at the time of testing. We selected this weight loss target after several pilot studies with the goal of achieving some consistency in the motivational states of the animals at different ages and not compromising health or activity. Mice are weighed before and after to food deprivation. To achieve the intended weight loss and motivational state, singly-housed adult mice were food deprived and habituated for 24 hours in the testing-cage. Water is available *ad libitum* at all times. The testing cage is an altered mouse cage (Thoren, #9) that has been equipped with an opaque 3D printed tunnel and lockable door. The custom designed tunnel (Meiccoms Engineering & Prototyping LLC, Spanish Fork, UT, USA) is 6cm in diameter, 18cm long, with stairs leading upward to an opening to the arena. The mice have access to explore the tunnel when the door is closed. Mice are housed in a room with a 10:00 – 22:00 dark cycle, so that testing is performed during the dark cycle. For testing, mice are moved into the behavior room prior to the start of testing for at least 30min for habituation to the new room. All testing is performed in the dark and video recording is done using infrared illumination and all manual procedures are done in the dark using red light. At the start of testing, the testing-cage is attached to the arena via the tunnel and the door is opened, allowing the mouse access to the arena, and video recording starts for the

Exploration phase. Mouse behavior is recorded continuously during the 30 min Exploration phase trial under infrared lights. Noldus Ethovision software v15 was used for video tracking. After completing the Exploration phase, the mouse is returned to the testing cage with water but no food until the Foraging phase 4 hours later. For the Foraging phase, the testing cage is again gently attached to the tunnel and access to the arena is possible and the Foraging trial begins. Video recording of the Foraging phase is performed for 30 minutes. After testing, mice are placed in their home cage with food and are returned to the mouse colony room. Between each Exploration and Foraging phase trial, the entire arena, including walls, platform, and tunnel are wiped clean with 70% ethanol.

***Preparation of sand and seed pots for the foraging assay***

Four white melamine pots (Carlisle S275, diameter 5.5cm, depth 4cm) were filled with 65g of sand. For the Exploration phase, one pot is filled with 50g of sand covered with 2.5g of seeds. On top of the seeds, a layer of 12g of sand is added to cover seeds. This sand is then covered with 0.5g of seeds. This pot is placed in position 2 in the arena. For the Foraging phase, one pot is filled with 50 g of sand, 3g of seeds on top of sand and additional 12g of sand to cover all seeds. This pot is placed in position 4 in the arena. All pots are weighed before and after the trial to measure the sand displaced from each pot. Remaining seeds and hulls left in the pot and on the platform are measured after each Exploration or Foraging trial to determine the amount of seeds consumed by the mouse during the trial. Used sand is collected after every trial and set aside. At the completion of all testing, the used sand is autoclaved before reuse in future trials.

***Foraging arena construction***

The assay cylinder walls were custom built with acrylic plastic (Delvies Plastics, Salt Lake City, UT, USA). The cylinder is made from a transparent acrylic tube and the 0.4 cm thick walls raise 50 cm above the platform, and were roughened with sandpaper to limit glaring and recording artifacts. The foraging platform sits on a supportive base and both are made from 0.5cm thick machine cut white matte ABS (Meiccoms Engineering & Prototyping LLC, Spanish Fork, UT, USA) with five 5.5cm diameter holes, one of which the 18cm long tunnel enters from underneath. The platform has a diameter of 35cm and sits 9.5 cm above the testing cage floor. The distance between the center of each 5.5cm hole pot is 16cm, and 12cm from center of platform.

***Arena tracking zones***

The arena is organized into zones that are used to breakdown the behavior and foraging strategies used by each animal in the assay. The arena is divided into five sectors and the boundary of each sector is the midpoint between two pots. The arena is further divided into three concentric circles, including the middle center zone, the intermediate zone and the outer wall zone. The outer radius of the *Intermediate* zone intersects with the center of the pots and tunnel entry. The radius of the *Center* zone is half the radius of the *Intermediate* zone. A zone is also created around each pot in the arena. *Pot* zones have a radius of 1.7x the radius of the pot itself. Finally, to learn about the behavior of the animal related to entries to and exits from the arena, we define zones around the tunnel entry. The *Tunnel Entry* zone aligns with the entry hole of the actual tunnel. The *In Tunnel* zone is covering the most peripheral area of the *Tunnel Entry* and tracks the mouse just before leaving the arena completely. Whenever the mouse is in the cage, the tracking system is recording the mouse as being in the *In Tunnel* zone. The *Tunnel Zone* area has the same radius as the *Pot* zones.

***Automated tracking***

At the start of the trial, the tracking begins with a 10 s delay to allow time for the testing-cage doors to be opened. The mouse is first tracked when it appears in the *In Tunnel zone* and position and movement is continuously recorded after this time point until the end of the 30 min Exploration or Foraging Phase. The XY position of the center of the mouse is video tracked with at a rate of 30 frames per second. Time spent in each zone, latency to visit a zone and number of visits to each zone, as well as the distance traveled, are calculated using the Ethovision software. The latency for the mouse to enter the arena is collected manually and defined as the time when the mouse has all four paws on the platform. Sand displacement and food consummation measures are collected and calculated manually.

### QUANTIFICATION AND STATISTICAL ANALYSIS

<u>Metabolic Phenotyping</u>

**CLAMS data graphing**

All CLAMS data from the torpor experiments were exported cropped to start at 6pm of the first day when the first light cycle starts and end at 12pm of the last day. Data was initially analyzed in CalR (*2*) and hourly average files were exported for further analysis in R. Body temperature collected through implants. The hourly average computed from CalR for each measure was visualized as line graphs with SEM using the plotly package in R.

**Modeling of CLAMS data**

For the statistical analysis of the torpor assay, we used the CLAMS data analyzed by CalR and Body temperature data and computed the average value in 6-hour time bins. Outliers in the data were managed as follows: Outlier values in Respiratory Exchange Ratio (RER) measures that are physiologically impossible (i.e. higher than 1.7) were set to 1. Outlier values in the food consumption data were removed by setting a maximum of 3.1 kcal/hr as this represents 1g of food consumed by a mouse in 1hr, which is the maximum food intake a mouse would reasonably consume. These outlier management approaches were guided by experts in CLAMS data analysis, including Dr. Amandine Chaix PhD and the University of Utah Metabolic Phenotyping Core facility.

To obtain single value measurements for each light or dark phase for each day for analysis, the two 6-hr measurements captured per phase were averaged, yielding a single mean value for each light-dark phase and day (Day.LD). Generalized linear modeling was performed on these values using a Gaussian distribution to test main genotype and interaction effects with a likelihood ratio test for full and nested models.

Mice were run in batches in our set of 8 CLAMS cages, typically aiming for a balance of 4 knockouts and 4 wild-type controls. Therefore, we included a term to absorb batch effect variability in our models.

We tested for a genotype effect using this model: value ~ Batch + Day.LD + Geno, during the total time, Pretorpor, Torpor and Refeeding phases.

We tested for Genotype x Day.LD interaction effects to identify effects specific to particular times of the assay using this model: value ~ Batch + Day.LD + Geno + Geno:Day.LD.

<u>Foraging Assay</u>

**Excursion Data Capture**

Our DeepFeats approach for analyzing modularity in foraging was performed as previously described (*1*, *3*), with modifications and advances. In our study, mice were tracked with Noldus Ethovision software. Noldus settings were used to define regions of interest in the foraging arena and indicated when the mouse was in each area. To ensure the tracking is equivalent across different mice, a Procrustes transformation of the XY coordinates was performed to put every tracking file in the same coordinate space. The track coordinates were zero'd to the center of the tunnel to the home cage. We then generated custom code in R to parse the raw Noldus tracking files into discrete, round trip home base excursions from the home cage tunnel. Each excursion is assigned a unique ID key that we call the Concise Idiosyncratic Module Alignment Report (CIMAR) string key. It stores the coordinates of the excursion in the data and the CIMAR string includes metadata regarding the mouse number, excursion number, sex, age, genotype, and phase. Next, custom code compares the CIMAR coordinates to the raw Noldus data files and constructs a new dataset that extracts 57 measures from the Noldus output, which we use for an initial statistical description of each excursion. The 57 measures are designed to capture a relatively comprehensive array of different behavioral and locomotor parameters, as well as describe interactions with food and non-food containing patches and exposed regions in the environment. These measures consist of shape, frequency, order and location statistics of an animal's X and Y movements, numbers of visits and time spent at different features in the arena, including food patches (Pots#2 and 4), non-food containing patches (Pots#1 and 3), the tunnel zone, wall zone and center zone of the arena and data describing locomotor patterns, including velocity, gait and distance traveled. The 57 measures for each excursion are scaled (normalized and centered across excursions) because they are in different units.

**Behavioral Measures to Resolve Modularity in Excursions**

A data matrix was constructed in which the rows are excursions performed by the mice, labeled by CIMAR keys, and the columns are the 57 behavioral measures. Dimension reduction was performed using Principal Component Analysis and the number of principal components (PCs) to retain for the identification of modules was defined based on the set that maximized cluster identification in the training dataset partition. Unsupervised clustering was performed on the retained 16 PCs. We used the Ward.D2 minimum variance method implemented using the "hclust" function in R to perform the clustering and define compact, spherical clusters. We then statistically define discrete excursion clusters from the results using the Dynamic Tree Cut algorithm (Langfelder et al., 2008). This is a powerful approach because it is adaptive to the shape of the dendrogram compared to typical constant height cutoff methods and offers the following advantages: (1) identification of nested clusters; (2) suitable for automation; and (3) can combine the advantages of hierarchical clustering and partitioning around medoids, giving better detection of outliers. We detect clusters using the “hybrid” method and use the DeepSplit parameter set to 4 and the minimum cluster size set to 20. The total number of clusters detected is quantified at each correlation threshold. Conceptually, more relaxed correlation threshold cutoffs could reduce cluster detection by retaining redundant measures that mask important effects from other measures. On the other hand, thresholds that are too stringent could reduce cluster detection by pruning informative measures. Our objective is to identify the threshold that uncovers the most informative and sensitive set of measures for resolving different clusters of excursions, setting the epoch for the discovery of potential modules.

**Statistical Validation of Significant Clusters of Excursions**
In our study, Dynamic Tree Cut will deeply cut branches in a dendrogram generating large numbers of small clusters if there are few bona fide relationships in the data. Thus, to test whether bona fide clusters of excursions exist in the data we implemented a random sampling procedure in R in which we randomly sample from the matrix of the retained behavioral measure data to break the relationships between the excursions and the measures. The sampled null data matrix is then subjected to the same clustering and quantification procedure to determine the number of clusters found by Dynamic Tree Cut (*4*). A null distribution is created from 10,000 iterations and compared to the observed number of clusters, which is expected to be significantly less than the null due to bona fide biological relationships between the excursions and set of retained measures. A lower tailed p value was computed to test this outcome.

**IGP Permutation Test for Stereotyped Behavioral Modules**
To test whether reproducible modules of behavior exist in the data for the foraging excursions, we use the in-group proportion (IGP) statistical method for testing for reproducible clusters between two datasets (*5*). We built a modified version of this function for parallelized computing to speed the analysis for large numbers of permutations. The excursion data for the mice is separated into a training data and test data partition for reproducibility testing. A balanced partition was generated according to genotype, sex and phase factors using the “createDataPartition” function in the caret package in R. Unsupervised hierarchical clustering was performed on the Training data partition excursions and 77 clusters were defined using Dynamic Tree Cut. Next, the centroids for each training data cluster were computed as the mean values of the behavioral data for the excursions in the cluster. The training data centroids were then used to compute the IGP statistic for each training data cluster based on the test partition data, thereby evaluating the reproducibility of each cluster.
As we previously described (*6*), we created a custom IGP permutation test that is based on a distance calculation, rather than the correlation implementation in the clusterRepro R package. The distance IGP testing framework was written in C++ and speeds the permutation test by many folds and is a more accurate replication of the clustering parameters used in the test data. We used this approach to compute p values for each cluster to determine whether the IGP value is greater than chance. False positives due to multiple testing errors were controlled using the q-value method (*7*). Modules are thus defined as significantly reproducible training partition

excursion clusters ($q < 0.1$). We detected 76 Modules in our dataset and one non-modular cluster. Each module detected was assigned an ID number and individual excursions in the data were annotated based on the module they match to. This approach facilitated quantifications of module expression frequency by the mice.

**Statistical Modeling of Module Expression Counts**
To statistically evaluate the genetic factors that significantly affect module expression frequency, we used generalized linear modeling functions implemented in R. The hypotheses tested are detailed in the text for each analysis and below. An interaction effect between module and genotype was tested using the model: Expression count ~ sex + age + genotype + module + genotype*module compared to a nested model of the main effects. Post-testing to define the specific modules that are affected was performed using a glm testing for a main effect of genotype on each individual module with the model: Expression count ~ sex + age + genotype. We filtered based on overall variance to remove modules with low expression variance across all mice in the study and reduce multiple testing errors, which is a proven two-step method for analyzing high dimensional data (*8*).

P-values were corrected for multiple testing errors by the q-value method to control the false discovery rate and pi0 was computed to determine true nulls in the data (*7*). Generalized linear modeling was performed using a Poisson distribution. We tested the goodness-of-fit of the Poisson for each module with a chi-square test of the residual deviance and degrees of freedom in R.

**Behavioral Cartography and Visualization**
To visualize the structure of the foraging behavior landscape and the relationships between modules and non-modular behavioral components, the raw data for 57 different behavioral measures was collected to describe each excursion for all 3928 excursions performed by all mice in the study. The data were analyzed and visualized using the PHATE algorithm (*9*, *10*) and annotated according to module and non-modular components as defined using the methods above.

**Statistical Analysis of Basic Foraging Measures using GLM**
We defined a set of 16 Basic “Keystone” measures of foraging that include, Distance Traveled, Relative Time in the Center (Time in the Center/Time on the Platform), Latency to Enter Arena, Latency to Visit Pot2 after Entering the Arena, Time on the Platform, Time at Pot2, Number of Sequences, Sand Displaced overall and at each Pot, Food Consumed, Relative Exposure (Time in the Center/ Time at the wall), Relative Latency to Enter the Arena (Exploration/Foraging), and Body Weight.

To statistically evaluate the variables that significantly affect these Keystone measures, we performed multiple regression analyses using R. Generalized linear modeling was performed using a Gamma(link=log) distribution. The hypotheses tested are detailed in the text for each analysis and below. An age effect was tested using the model: Feature expression ~ Phase + Sex + Genotype + Age. A genotype effect was tested using the model: Feature expression ~ Phase + Sex + Age + Genotype. An age by genotype interaction was tested using the model: Feature expression ~ Phase + Sex + Age + Genotype + Age*Genotype. A sex by genotype interaction was tested using the model: Feature expression ~ Age + Phase + Sex + Genotype + Sex*Genotype. An age by sex by genotype interaction was tested using the model: Feature expression ~ Age + Phase + Sex + Genotype + Age*Sex*Genotype. An age by phase by genotype interaction was tested using the model: Feature expression ~ Age + Phase + Sex + Genotype + Age*Phase*Genotype. Posttests were calculated using a T-test implemented in R. Each Keystone Feature was tested for a significant genotype effect (p<0.05) for each pair (KO vs WT).

The time spent at Pot2 during the 30 min trials was compared between wt and ko mice using a mixed model approach as follows:

For an Interval (1min time bins) by Age by Genoype interaction:
```
model0 <- lmer(Time ~ Sex + Geno + Interval + AgeGroup + Interval:AgeGroup:Geno + (1|Mouse)
model1 <- lmer(Time ~ Sex + Geno + Interval + AgeGroup + (1|Mouse)
```

For a genotype effect:
```
model0 <- lmer(Time ~ Sex + Interval + AgeGroup + Geno + (1|Mouse)
model1 <- lmer(Time ~ Sex + Interval + AgeGroup + (1|Mouse)
```

For an  Age by Geno effect:
```
model0 <- lmer(Time ~ Sex + Interval + AgeGroup + Geno + AgeGroup:Geno + (1|Mouse)
model1 <- lmer(Time ~ Sex + Interval + AgeGroup + Geno + (1|Mouse)
```

For a Sex by Genotype effect:
```
model0 <- lmer(Time ~ Sex + Interval + AgeGroup + Geno + Sex:Geno + (1|Mouse)
model1 <- lmer(Time ~ Sex + Interval + AgeGroup + Geno + (1|Mouse)
```

For an Interval by Genotype  effect:
```
model0 <- lmer(Time ~ Sex + Interval + AgeGroup + Geno + Interval:Geno + (1|Mouse)
model1 <- lmer(Time ~ Sex + Interval + AgeGroup + Geno + (1|Mouse)
```

For an Age by Sex effect:
```
model0 <- lmer(Time ~ Sex + Interval + AgeGroup + Geno + AgeGroup:Sex + (1|Mouse)
model1 <- lmer(Time ~ Sex + Interval + AgeGroup + Geno + (1|Mouse)
```

For an  Age  effect:
```
model0 <- lmer(Time ~ Sex + Interval + Geno + AgeGroup + (1|Mouse)
model1 <- lmer(Time ~ Sex + Interval + Geno + (1|Mouse)
```

**PCA Regression Analysis of Full Foraging Profiles (Basic + Module Measures)**

Separately for Exploration and Foraging phase data, we performed PCA on the centered and scaled foraging profiles (93 features total, including all basic Keystone measures and module expression count data) of all 156 mice. Within each phase, we separated young (n=43) and aged (n=113) mice and performed logistic regression to predict the binary genotype (encoded as 0 and 1) from the principal component features. We found a local peak in the adjusted r-squared statistic at 7 PCs for aged mice in the foraging phase.

**SOFTWARE USED**

Rstudio
Python
Noldus Ethovision
Prism
Dynamic Tree Cut (Bioconductor)
PHATE (CRAN)

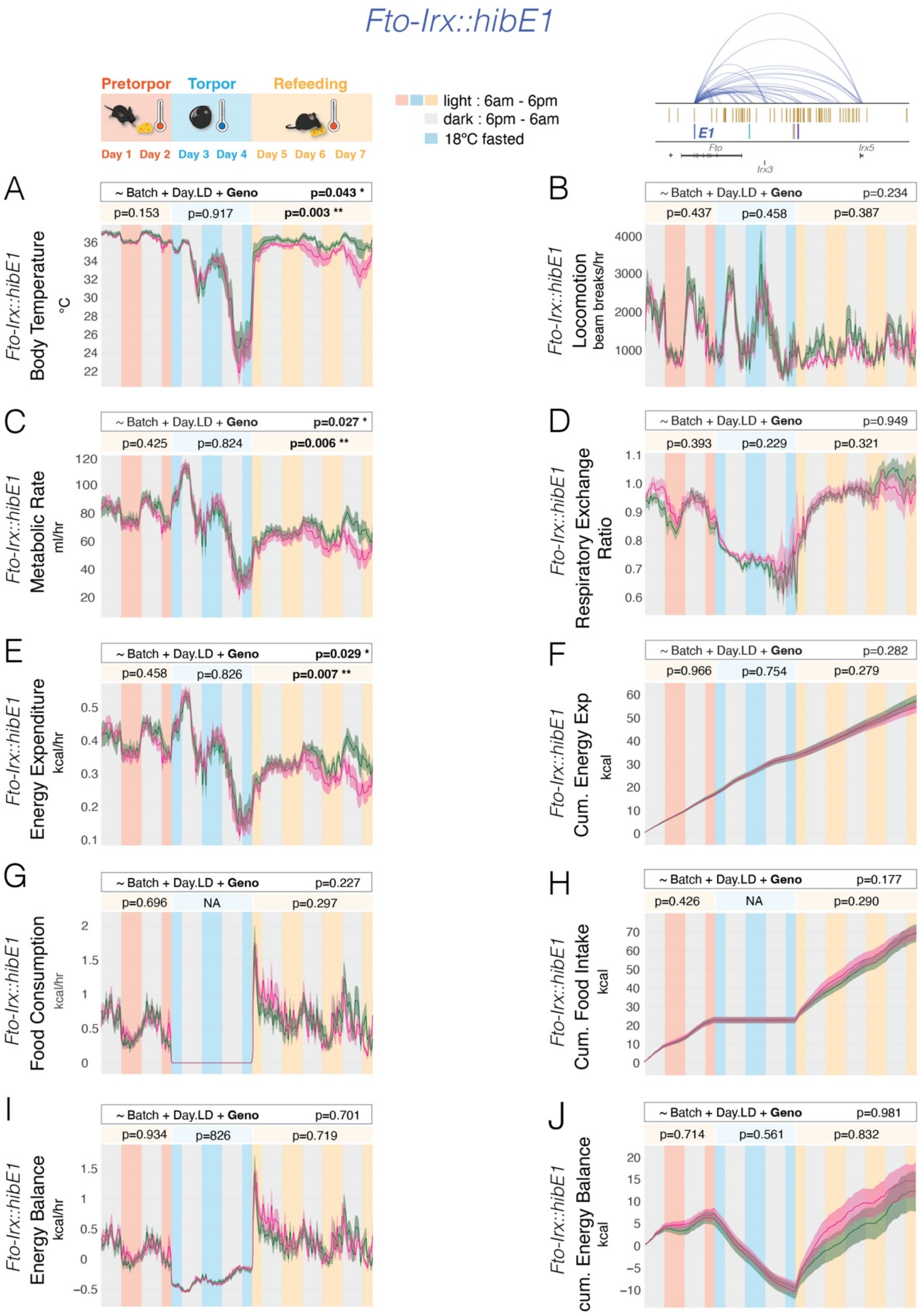
Fto-Irx::hibE1
Pretorpor
Torpor
Refeeding
Day 1
Day 2
Day 3
Day 4
Day 5
Day 6
Day 7
light : 6am - 6pm
dark : 6pm - 6am
18°C fasted
E1
Fto
Irx3
Irx5
A
~ Batch + Day.LD + Geno p=0.043 *
p=0.153 p=0.917 p=0.003 **
Fto-Irx::hibE1
Body Temperature
°C
B
~ Batch + Day.LD + Geno p=0.234
p=0.437 p=0.458 p=0.387
Fto-Irx::hibE1
Locomotion
beam breaks/hr
C
~ Batch + Day.LD + Geno p=0.027 *
p=0.425 p=0.824 p=0.006 **
Fto-Irx::hibE1
Metabolic Rate
ml/hr
D
~ Batch + Day.LD + Geno p=0.949
p=0.393 p=0.229 p=0.321
Fto-Irx::hibE1
Respiratory Exchange Ratio
E
~ Batch + Day.LD + Geno p=0.029 *
p=0.458 p=0.826 p=0.007 **
Fto-Irx::hibE1
Energy Expenditure
kcal/hr
F
~ Batch + Day.LD + Geno p=0.282
p=0.966 p=0.754 p=0.279
Fto-Irx::hibE1
Cum. Energy Exp
kcal
G
~ Batch + Day.LD + Geno p=0.227
p=0.696 NA p=0.297
Fto-Irx::hibE1
Food Consumption
kcal/hr
H
~ Batch + Day.LD + Geno p=0.177
p=0.426 NA p=0.290
Fto-Irx::hibE1
Cum. Food Intake
kcal
I
~ Batch + Day.LD + Geno p=0.701
p=0.934 p=826 p=0.719
Fto-Irx::hibE1
Energy Balance
kcal/hr
J
~ Batch + Day.LD + Geno p=0.981
p=0.714 p=0.561 p=0.832
Fto-Irx::hibE1
cum. Energy Balance
kcal

**Fig. S1. Metabolic Profiles and Body Composition in *Fto-Irx::hibE1*$^{-/-}$ Mice Across Nutritional States Mimicking Hibernation Cycles reveal Decreased Metabolic Rate and Energy Expenditure during Refeeding Phase.**

(**A-J**) The plots show $Tb_i$ and CLAMS metabolic measures for *Fto-Irx::hibE1* $^{-/-}$ (pink, n=14) versus $^{+/+}$ (green, n=11) adult female littermates in pre-torpor (ad libitum food + 25°C), torpor (48 hrs FD + 18°C), and refeeding phases (ad libitum food + 25°C). Significant differences between *Fto-Irx::hibE1* $^{-/-}$ and $^{+/+}$ mice were observed in a regression analysis for a main effect of genotype involving decreased $Tb_i$ (A), metabolic rate (C, MR), and energy expenditure (E, EE) in the knockouts in the refeeding phase. Dark pink and green lines show the mean, while shaded areas show S.E.M. The genotype effect p-value for the entire assay is shown on top, while p-values for each phase are below. *p<0.05, **p<0.01, ***p<0.001

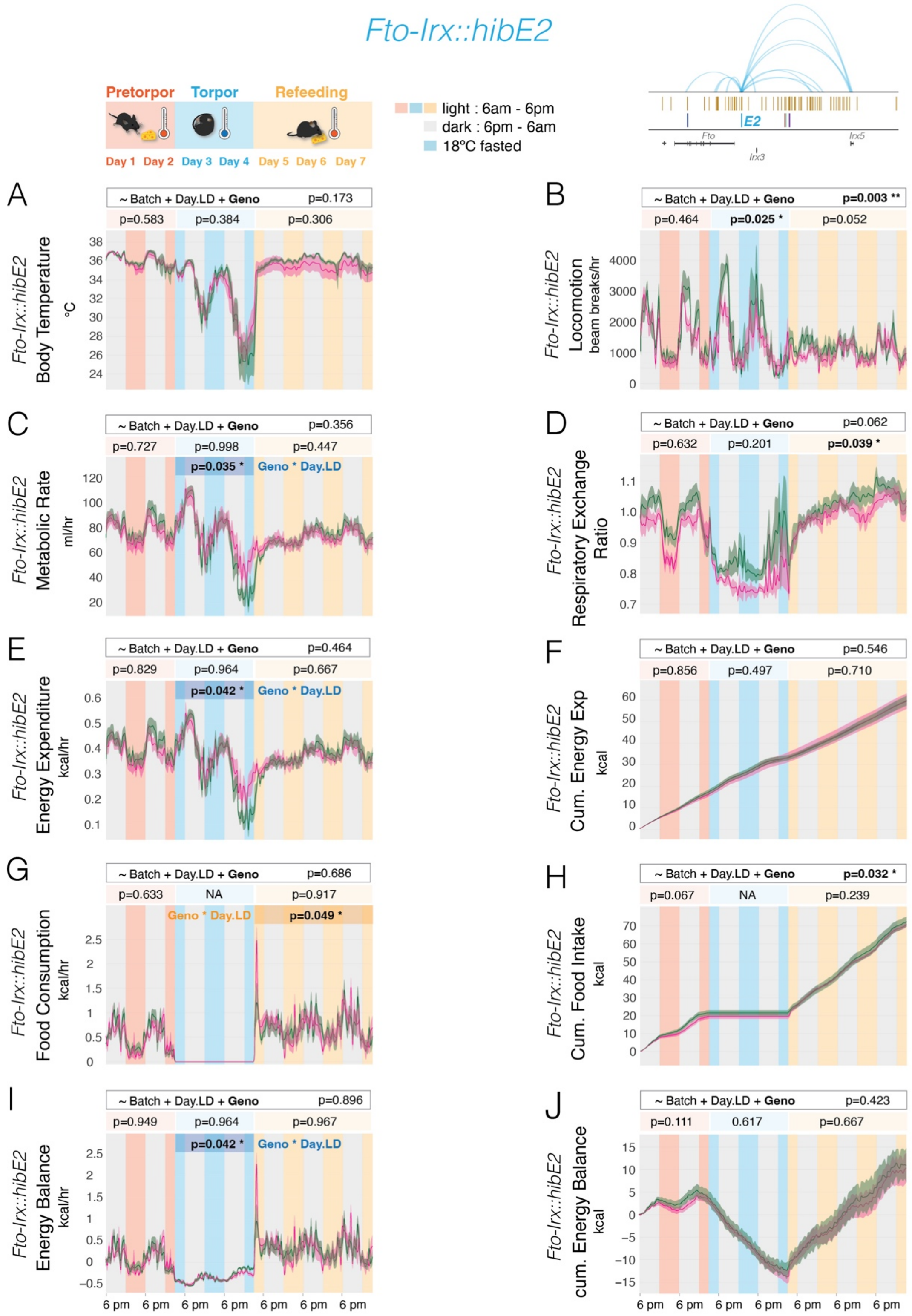

Fto-Irx::hibE2
Pretorpor
Torpor
Refeeding
Day 1 Day 2 Day 3 Day 4 Day 5 Day 6 Day 7
light : 6am - 6pm
dark : 6pm - 6am
18°C fasted
E2
Fto
Irx3
Irx5
A
~ Batch + Day.LD + Geno p=0.173
p=0.583 p=0.384 p=0.306
Fto-Irx::hibE2 Body Temperature °C
B
~ Batch + Day.LD + Geno p=0.003 **
p=0.464 p=0.025 * p=0.052
Fto-Irx::hibE2 Locomotion beam breaks/hr
C
~ Batch + Day.LD + Geno p=0.356
p=0.727 p=0.998 p=0.447
p=0.035 * Geno * Day.LD
Fto-Irx::hibE2 Metabolic Rate ml/hr
D
~ Batch + Day.LD + Geno p=0.062
p=0.632 p=0.201 p=0.039 *
Fto-Irx::hibE2 Respiratory Exchange Ratio
E
~ Batch + Day.LD + Geno p=0.464
p=0.829 p=0.964 p=0.667
p=0.042 * Geno * Day.LD
Fto-Irx::hibE2 Energy Expenditure kcal/hr
F
~ Batch + Day.LD + Geno p=0.546
p=0.856 p=0.497 p=0.710
Fto-Irx::hibE2 Cum. Energy Exp kcal
G
~ Batch + Day.LD + Geno p=0.686
p=0.633 NA p=0.917
Geno * Day.LD p=0.049 *
Fto-Irx::hibE2 Food Consumption kcal/hr
H
~ Batch + Day.LD + Geno p=0.032 *
p=0.067 NA p=0.239
Fto-Irx::hibE2 Cum. Food Intake kcal
I
~ Batch + Day.LD + Geno p=0.896
p=0.949 p=0.964 p=0.967
p=0.042 * Geno * Day.LD
Fto-Irx::hibE2 Energy Balance kcal/hr
J
~ Batch + Day.LD + Geno p=0.423
p=0.111 0.617 p=0.667
Fto-Irx::hibE2 cum. Energy Balance kcal
6 pm 6 pm 6 pm 6 pm 6 pm 6 pm 6 pm

**Fig. S2. Metabolic Profiles and Body Composition in *Fto-Irx::hibE2*$^{-/-}$ Mice Across Nutritional States Mimicking Hibernation Cycles reveal Elevated Metabolic Rate and Decreased Locomotion During Torpor Phase.**

(**A-J**) The plots show $Tb_i$ and CLAMS metabolic measures for *Fto-Irx::hibE2* $^{-/-}$ (pink, n=7) versus $^{+/+}$ (green, n=8) adult female littermates in pre-torpor (ad libitum food + 25°C), torpor (48 hrs FD + 18°C), and refeeding phases (ad libitum food + 25°C). Significant differences between *Fto-Irx::hibE2* $^{-/-}$ and $^{+/+}$ mice were observed in the torpor phase for an interaction effect between genotype x Day.LD for metabolic rate (C, MR), energy expenditure (E, EE), and energy balance (I, EB), as well as in the refeeding phase for Food Consumption (G). The phenotypic effect involves increases in the knockout mice near the end of the torpor phase. Significant main effects of genotype were observed for locomotion in the torpor phase (B), respiratory exchange ratio in the refeeding phase (D), and over the entire assay for Cumulative food intake (H). Dark pink and green lines show the mean, while shaded areas show S.E.M. The genotype effect p-value for the entire assay is shown on top, while p-values for each phase are below. Significant interactions between genotype X Day.LD are shown in blue (torpor phase) and orange (refeeding phase). *p<0.05, **p<0.01, ***p<0.001

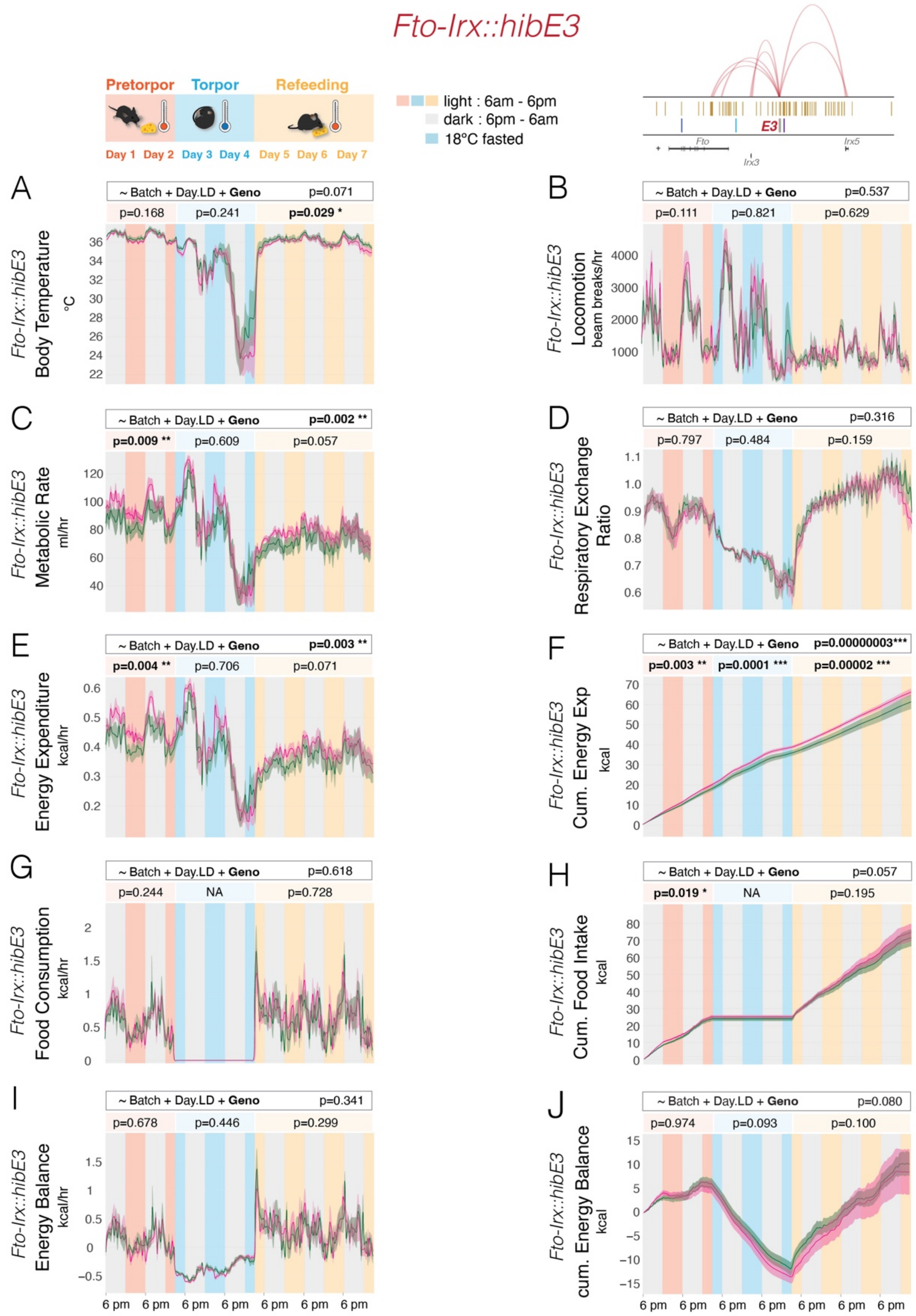

Fto-Irx::hibE3
Pretorpor
Torpor
Refeeding
Day 1 Day 2 Day 3 Day 4 Day 5 Day 6 Day 7
light : 6am - 6pm
dark : 6pm - 6am
18°C fasted
E3
Fto
Irx3
Irx5
A
~ Batch + Day.LD + Geno p=0.071
p=0.168 p=0.241 p=0.029 *
Fto-Irx::hibE3 Body Temperature °C
B
~ Batch + Day.LD + Geno p=0.537
p=0.111 p=0.821 p=0.629
Fto-Irx::hibE3 Locomotion beam breaks/hr
C
~ Batch + Day.LD + Geno p=0.002 **
p=0.009 ** p=0.609 p=0.057
Fto-Irx::hibE3 Metabolic Rate ml/hr
D
~ Batch + Day.LD + Geno p=0.316
p=0.797 p=0.484 p=0.159
Fto-Irx::hibE3 Respiratory Exchange Ratio
E
~ Batch + Day.LD + Geno p=0.003 **
p=0.004 ** p=0.706 p=0.071
Fto-Irx::hibE3 Energy Expenditure kcal/hr
F
~ Batch + Day.LD + Geno p=0.00000003***
p=0.003 ** p=0.0001 *** p=0.00002 ***
Fto-Irx::hibE3 Cum. Energy Exp kcal
G
~ Batch + Day.LD + Geno p=0.618
p=0.244 NA p=0.728
Fto-Irx::hibE3 Food Consumption kcal/hr
H
~ Batch + Day.LD + Geno p=0.057
p=0.019 * NA p=0.195
Fto-Irx::hibE3 Cum. Food Intake kcal
I
~ Batch + Day.LD + Geno p=0.341
p=0.678 p=0.446 p=0.299
Fto-Irx::hibE3 Energy Balance kcal/hr
J
~ Batch + Day.LD + Geno p=0.080
p=0.974 p=0.093 p=0.100
Fto-Irx::hibE3 cum. Energy Balance kcal
6 pm 6 pm 6 pm 6 pm 6 pm 6 pm 6 pm

**Fig. S3. Metabolic Profiles and Body Composition in *Fto-Irx::hibE3*$^{-/-}$ Mice Across Nutritional States Mimicking Hibernation Cycles reveal increased Metabolic Rate and Energy Expenditure.**

(**A-J**) The plots show $Tb_i$ and CLAMS metabolic measures for *Fto-Irx::hibE3* $^{-/-}$ (pink, n=7) versus $^{+/+}$ (green, n=6) adult female littermates in pre-torpor (ad libitum food + 25°C), torpor (48 hrs FD + 18°C), and refeeding phases (ad libitum food + 25°C). Significant differences between *Fto-Irx::hibE3* $^{-/-}$ and $^{+/+}$ mice were observed in a regression analysis for a genotype effect for metabolic rate (C, MR) and energy expenditure (E, EE) that involves increases across the whole assay and the pre-torpor phase. Cumulative EE was significantly increased in the knockouts in all phases (F, Cum EE). A significant increase in cumulative food intake in the pre-torpor phase was also observed (H). A significant decrease in $Tb_i$ was found in the refeeding phase along with a trend for decreased $Tb_i$ across the whole assay (A). Dark pink and green lines show the mean, while shaded areas show S.E.M. The genotype effect p-value for the entire assay is shown on top, while p-values for each phase are below. *$p<0.05$, **$p<0.01$, ***$p<0.001$

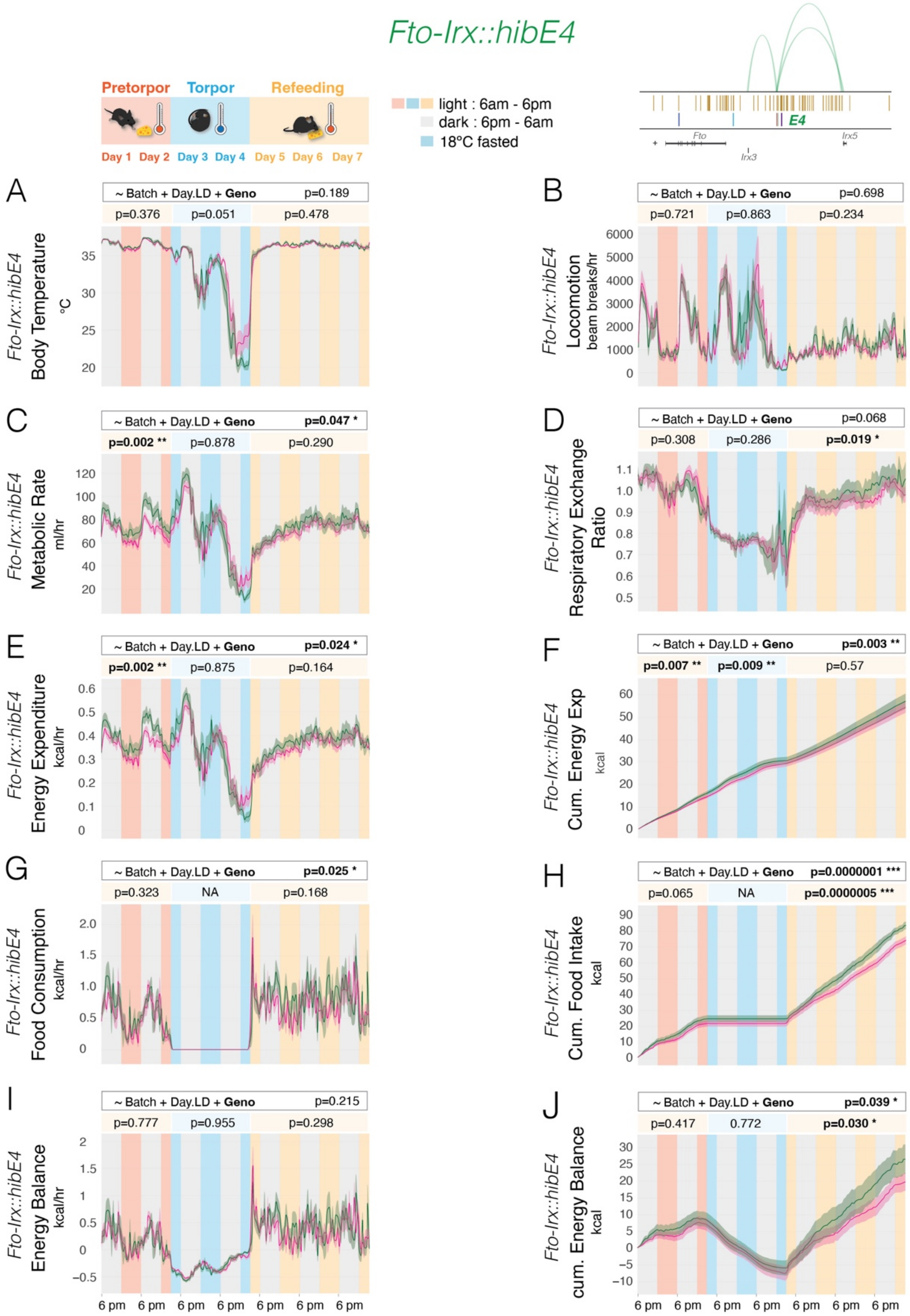

Fto-Irx::hibE4
Pretorpor
Torpor
Refeeding
Day 1 Day 2 Day 3 Day 4 Day 5 Day 6 Day 7
light : 6am - 6pm
dark : 6pm - 6am
18°C fasted
E4
Fto
Irx3
Irx5
A
~ Batch + Day.LD + Geno p=0.189
p=0.376 p=0.051 p=0.478
Fto-Irx::hibE4 Body Temperature °C
B
~ Batch + Day.LD + Geno p=0.698
p=0.721 p=0.863 p=0.234
Fto-Irx::hibE4 Locomotion beam breaks/hr
C
~ Batch + Day.LD + Geno p=0.047 *
p=0.002 ** p=0.878 p=0.290
Fto-Irx::hibE4 Metabolic Rate ml/hr
D
~ Batch + Day.LD + Geno p=0.068
p=0.308 p=0.286 p=0.019 *
Fto-Irx::hibE4 Respiratory Exchange Ratio
E
~ Batch + Day.LD + Geno p=0.024 *
p=0.002 ** p=0.875 p=0.164
Fto-Irx::hibE4 Energy Expenditure kcal/hr
F
~ Batch + Day.LD + Geno p=0.003 **
p=0.007 ** p=0.009 ** p=0.57
Fto-Irx::hibE4 Cum. Energy Exp kcal
G
~ Batch + Day.LD + Geno p=0.025 *
p=0.323 NA p=0.168
Fto-Irx::hibE4 Food Consumption kcal/hr
H
~ Batch + Day.LD + Geno p=0.0000001 ***
p=0.065 NA p=0.0000005 ***
Fto-Irx::hibE4 Cum. Food Intake kcal
I
~ Batch + Day.LD + Geno p=0.215
p=0.777 p=0.955 p=0.298
Fto-Irx::hibE4 Energy Balance kcal/hr
J
~ Batch + Day.LD + Geno p=0.039 *
p=0.417 0.772 p=0.030 *
Fto-Irx::hibE4 cum. Energy Balance kcal
6 pm 6 pm 6 pm 6 pm 6 pm 6 pm 6 pm

**Fig. S4. Metabolic Profiles and Body Composition in *Fto-Irx::hibE4$^{-/-}$* Mice Across Nutritional States Mimicking Hibernation Cycles reveal decreased Metabolic Rate, Energy Expenditure, Locomotion, and Energy Balance.**

(**A-J**) The plots show $Tb_i$ and CLAMS metabolic measures for *Fto-Irx::hibE4* $^{-/-}$ (pink, n=8) versus $^{+/+}$ (green, n=6) adult female littermates in pre-torpor (ad libitum food + 25°C), torpor (48 hrs FD + 18°C), and refeeding phases (ad libitum food + 25°C). Significant differences between *Fto-Irx::hibE4* $^{-/-}$ and $^{+/+}$ mice were observed in a regression analysis for a genotype effect for metabolic rate (C, MR) and energy expenditure (E, EE) that involves a decrease in the pre-torpor phase. Food consumption (G) was significantly decreased over the whole assay. Respiratory exchange ratio (C) was decreased during the refeeding phase. Cumulative EE (F), cumulative food intake (H), and cumulative energy balance (J) was significantly decreased in the knockouts. Dark pink and green lines show the mean, while shaded areas show S.E.M. The genotype effect p-value for the entire assay is shown on top, while p-values for each phase are below. *$p<0.05$, **$p<0.01$, ***$p<0.001$

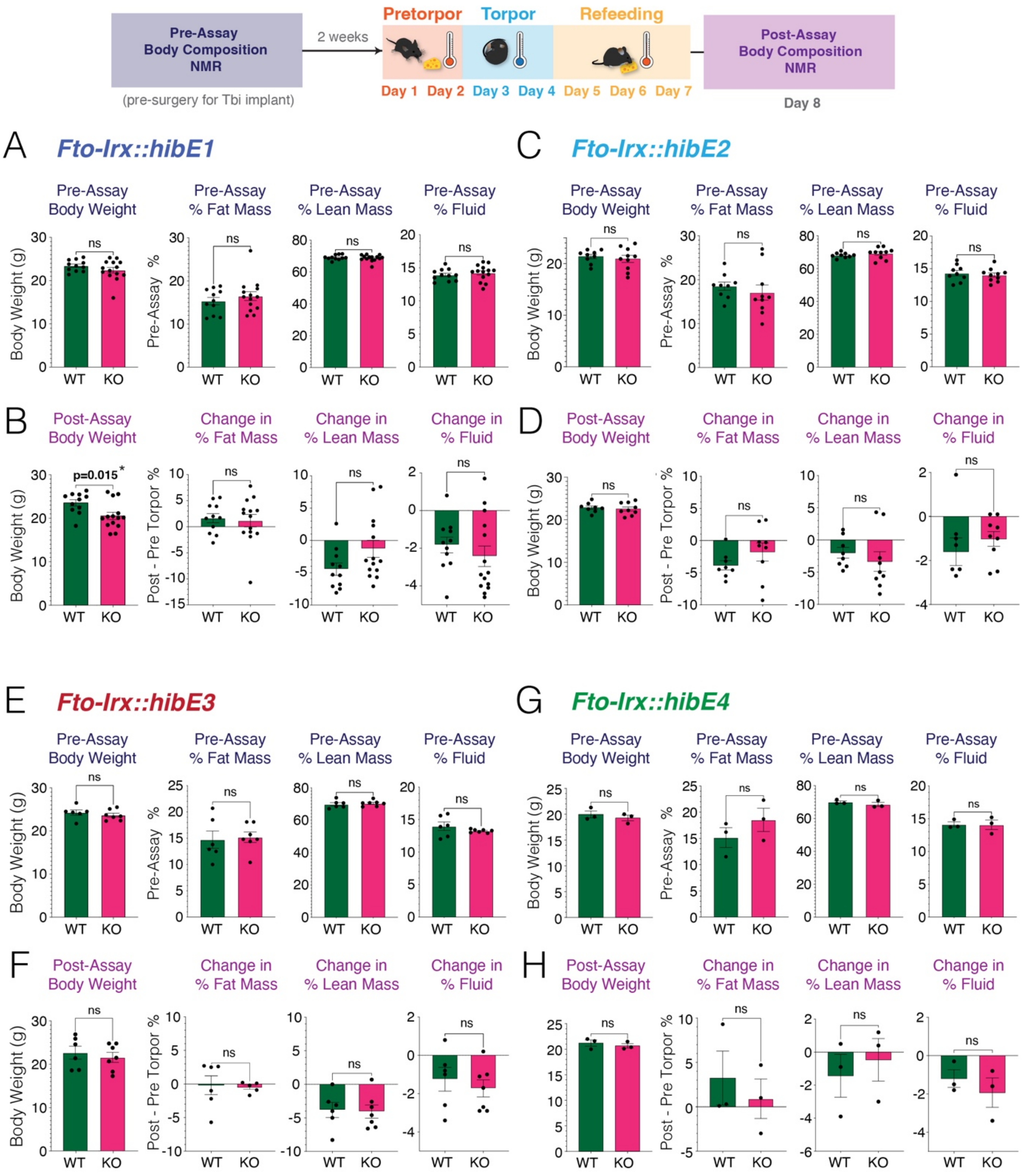

Pre-Assay
Body Composition
NMR
(pre-surgery for Tbi implant)
2 weeks
Pretorpor
Torpor
Refeeding
Day 1
Day 2
Day 3
Day 4
Day 5
Day 6
Day 7
Post-Assay
Body Composition
NMR
Day 8
A
Fto-Irx::hibE1
B
C
Fto-Irx::hibE2
D
E
Fto-Irx::hibE3
F
G
Fto-Irx::hibE4
H
Pre-Assay Body Weight
Pre-Assay % Fat Mass
Pre-Assay % Lean Mass
Pre-Assay % Fluid
Post-Assay Body Weight
Change in % Fat Mass
Change in % Lean Mass
Change in % Fluid
Body Weight (g)
Pre-Assay %
Post - Pre Torpor %
p=0.015*
ns
WT
KO

**Fig. S5. Body Weight and Composition in *Fto-Irx::hibE cis*-element Knockout Mice Before and After Profiling in the Torpor Metabolic Assay.**

(**A-H**) The barplots show the pre-assay (A, C, E, G; dark blue titles) and post-assay (B, D, F, H; purple titles) body weight and body composition data for the CLAMS metabolic torpor assay (see top schematic) for $^{-/-}$ versus $^{+/+}$ adult female littermates for the *Fto-Irx::hibE1* (A,B; n=11-14)*, Fto-Irx::hibE2* (C,D; n=7-8)*, Fto-Irx::hibE3* (E,F; n=6-7)*, Fto-Irx::hibE4* (G,H; n=3) mouse lines. *Fto-Irx::hibE1*-/- show significantly reduced body weight post-assay (B). Significant effects were not observed for other measures or mouse lines. Changes in % fat mass, lean mass or fluid (purple text measures) were computed from the difference in post-assay % minus pre-assay % values. Statistical analysis was performed using an unpaired two-tailed t-test to compare $^{-/-}$ to $^{+/+}$ mice. ***p<0.0001, **p<0.01, *p<0.05.

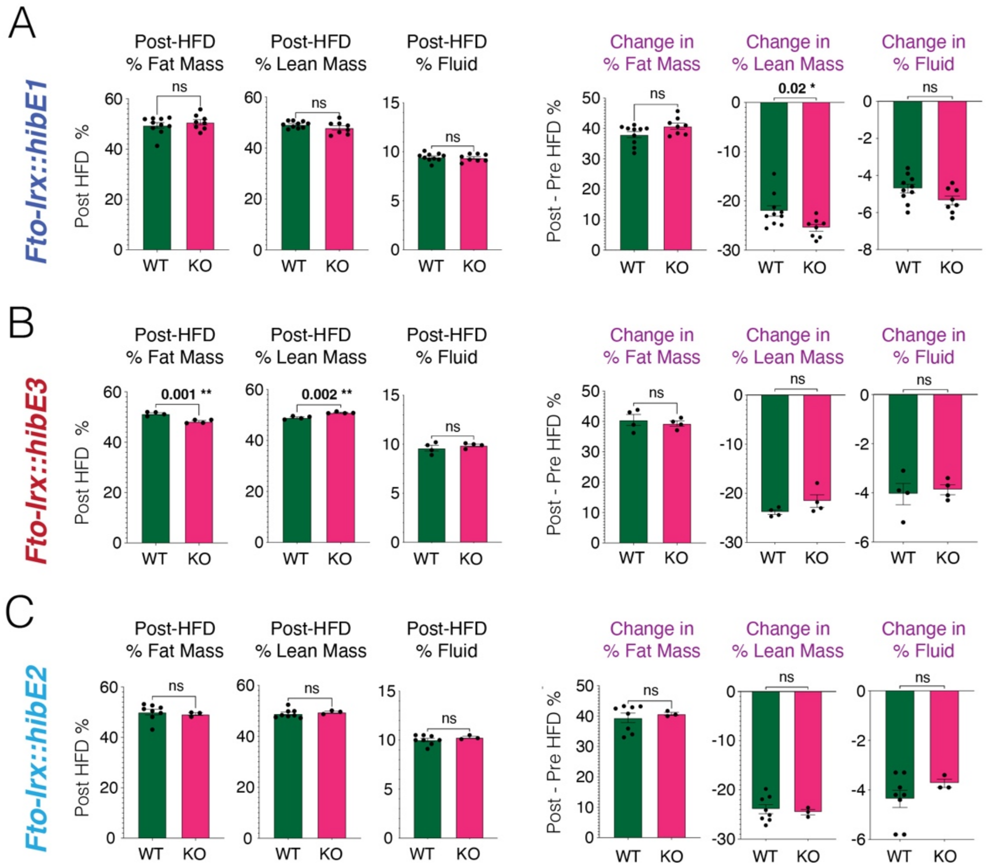
A
Fto-Irx::hibE1
Post-HFD % Fat Mass
Post-HFD % Lean Mass
Post-HFD % Fluid
Change in % Fat Mass
Change in % Lean Mass
Change in % Fluid
Post HFD %
Post - Pre HFD %
ns
0.02 *
WT
KO
B
Fto-Irx::hibE3
0.001 **
0.002 **
C
Fto-Irx::hibE2

**Fig. S6. Body Composition for *Fto-Irx::hibE cis*-element Knockout Mice on a High Fat Diet reveals opposing lean mass phenotypes for *Fto-Irx::hibE1* versus *Fto-Irx::hibE3* Knockouts.**

(**A-C**) The barplots show the body composition data for adult male mice placed on a high fat diet (HFD) for 22 weeks in a comparison of $^{-/-}$ (pink) versus $^{+/+}$ (green) littermates for the *Fto-Irx::hibE1* (A; -/- n=8; +/+ n=10), *Fto-Irx::hibE3* (B; -/- n=4; +/+ n=4), and *Fto-Irx::hibE2* (C; -/- n=3; +/+ n=8) mouse lines. *Fto-Irx::hibE1*-/- mice show a significantly greater reduction in the % of body composition that is lean mass at the end of HF diet relative to the start of the diet (A, Change in % Lean Mass, purple text). *Fto-Irx::hibE3*-/- mice show significantly reduced % fat mass at the end of the HFD (B, Post-HFD % Fat Mass) and increased % lean mass (B, Post-HFD % Lean Mass). Significant effects were not observed for the *Fto-Irx::hibE2*-/- mice. Changes in % fat mass, lean mass, or fluid (purple text measures) were computed from the difference in post-HFD % minus pre-HFD % values. Unpaired two-tailed t-test comparing $^{-/-}$ to $^{+/+}$ mice. WT, +/+; KO, -/-. ***p<0.0001, **p<0.01, *p<0.05.

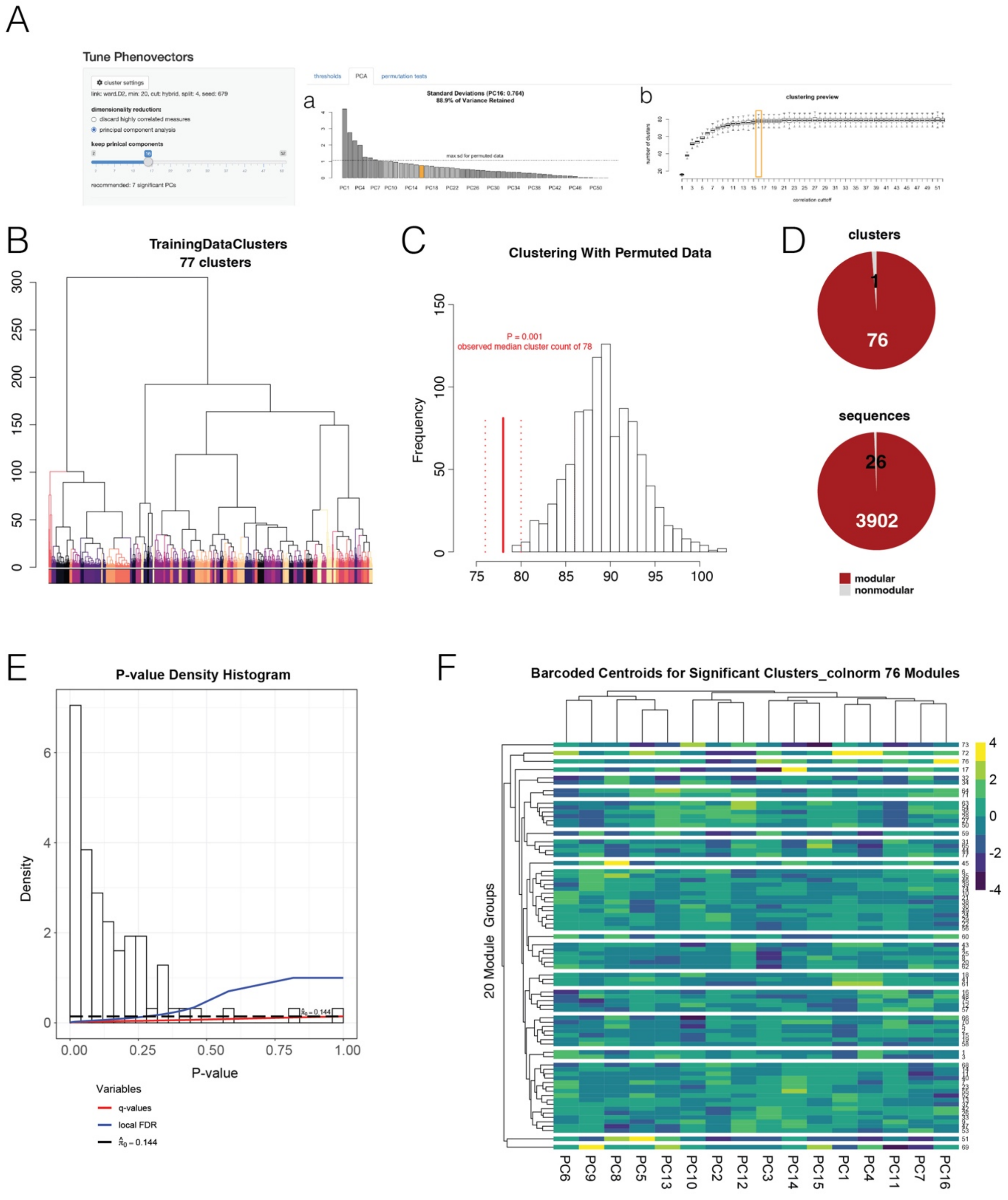

A
Tune Phenovectors
a
Standard Deviations (PC16: 0.764)
88.9% of Variance Retained
b
clustering preview
B
TrainingDataClusters
77 clusters
C
Clustering With Permuted Data
P = 0.001
observed median cluster count of 78
Frequency
D
clusters
1
76
sequences
26
3902
modular
nonmodular
E
P-value Density Histogram
Density
P-value
Variables
q-values
local FDR
F
Barcoded Centroids for Significant Clusters_colnorm 76 Modules
20 Module Groups
PC6 PC9 PC8 PC5 PC13 PC10 PC2 PC12 PC3 PC14 PC15 PC1 PC4 PC11 PC7 PC16

**Fig. S7. Identification of foraging modules in *Fto-Irx::hibE3* mice.**

(**A**) The PCA Variance Retained plot (a) shows the number of principle components (PCs) and proportion of variance explained by each PC in our DeepFeats algorithm. The chosen PC cutoff is in orange at the elbow of the plot and captures 88.9% of variance in the data for hundreds of measures describing discrete round trip foraging excursions from the home. The Clustering Preview (b) plot shows the number of clusters discovered with the selected PCs based on different cutoffs for the number of PCs chosen. The data reveal that the maximum cluster number and minimum standard deviation in terms of cluster number is achieved using 16 PCs.
(**B**) Dynamic tree cutting yielded 77 candidate clusters of excursions in the training data. This training data clusters were tested for significant reproducibility in the test data partition to uncover significant modules.
(**C**) The plot shows the results of a permutation test in which the data describing each excursion is randomly permuted and the number of excursion clusters uncovered by DeepFeats from permuted data is shown in the histogram and compared to the observed number (red). The results show significantly fewer clusters are identified in the true data compared to permuted data, consistent with a bona fide clusters compared to random.
(**D and E**) The pie charts show the number of training data clusters that are significant modules showing reproducibility in the test data (red) versus the number that are not significantly reproducible in the test data (grey) (ie. nonmodular) (D). The second pie chart below shows the number of round-trip excursions found to be modular (red) versus non-modular (grey) (D). An in-group proportion permutation test was performed to define clusters in the training data that are reproducible in the test data, (Q-value <0.1 for significant modules). The histogram shows the p-values from the IGP test for q-value calculation (E).

(**F**) The heatmap shows the values for the PCs for the centroids for each significant excursion cluster (module) scaled by column. The results show how different PCs summarizing the 52 behavioral measures differentiate each module and unsupervised hierarchical clustering on the centroids shows groups of similar modules.

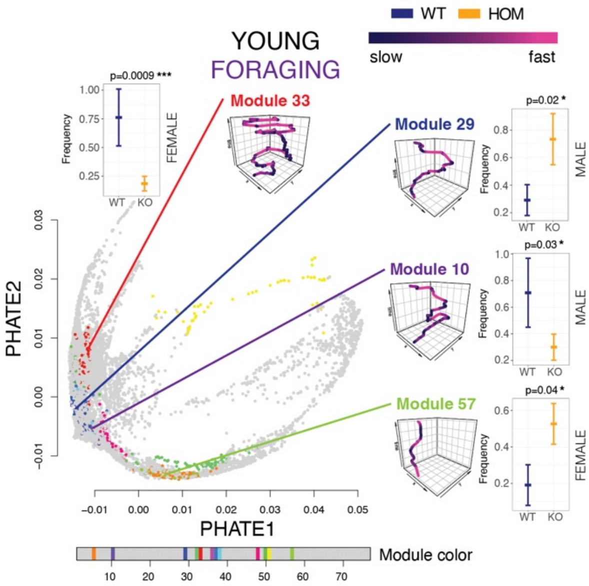

WT
HOM
YOUNG
FORAGING
slow
fast
Module 33
Module 29
Module 10
Module 57
p=0.0009 ***
FEMALE
p=0.02 *
MALE
p=0.03 *
MALE
p=0.04 *
FEMALE
Frequency
WT
KO
PHATE1
PHATE2
Module color

**Fig. S8. Significantly affected foraging modules in young adult *Fto-Irx::hibE3*$^{-/-}$ mice.**

The PHATE map plots show the modules and excursions that are significantly differentially expressed in $^{-/-}$ versus $^{+/+}$ mice in young adults in the Foraging phase. The purple traces show foraging movement patterns representative of each affected example module shaded by velocity. Regression analysis revealed significant effects of genotype on the expression of these modules and the plots show the expression frequency of each foraging module in $^{+/+}$ versus $^{-/-}$ mice. Line plots show the mean and bars show SEM for the module expression frequency per mouse in *Fto-Irx::hibE3* WT (+/+, blue) and KO (-/-, orange) male or female mice (as indicated). See table in Figure 4 for number of mice per group. ***p<0.0001, **p<0.01, *p<0.05.

### SUPPLEMENTAL MOVIES

**Movie S1. Second-guessing behavior during a naturalistic foraging assay.** The video shows footage of a mouse foraging in the Foraging phase of our assay. Despite food in the new food patch (Pot4), the mouse repeatedly visits and digs in the former food patch (pot2), which was learned to contain food in the naïve Exploration phase. We refer to this perseverative revisiting of the former food patch as "second-guessing".

**Movie S2. The video shows the 3D projected PHATE map of all mouse excursions and foraging module classifications of each excursion.** Dots represent individual foraging excursions described by 52 measures and embedded in 3-D space. Excursions are colored by classifications to modules based on DeepFeats analysis and the data show that excursions assigned to the same module are near in space and therefore similar, as expected.

**Movie S3. The video shows the 3D projected PHATE map of all mouse excursions for modules showing a significant difference in expression between young adult *Fto-Irx::hibE3* $^{-/-}$ versus $^{+/+}$ mice in the Foraging phase.** The results show that a small subset of modules are significantly affected (colored dots; colored by module type) in the young adult *Fto-Irx::hibE3*$^{-/-}$ mice compared to aged -/- mice (Video S3).

**Movie S4. The video shows the 3D projected PHATE map of all mouse excursions for modules showing a significant difference in expression between aged *Fto-Irx::hibE3* $^{-/-}$ versus $^{+/+}$ mice in the Foraging phase.** The results show that a specific subset of modules are significantly affected (colored dots; colored by module type) in aged *Fto-Irx::hibE3*$^{-/-}$ mice, and many excursions are not significantly affected (grey dots).
